# Pairing to the microRNA 3′ region occurs through two alternative binding modes, with affinity shaped by nucleotide identity as well as pairing position

**DOI:** 10.1101/2021.04.13.439700

**Authors:** Sean E. McGeary, Namita Bisaria, David P. Bartel

**Author notes:** These authors contributed equally to this work (Sean E. McGeary, Namita Bisaria).

## Abstract

MicroRNAs (miRNAs), in association with Argonaute (AGO) proteins, direct repression by pairing to sites within mRNAs. Compared to pairing preferences of the miRNA seed region (nucleotides 2–8), preferences of the miRNA 3′ region are poorly understood, due to the sparsity of measured affinities for the many pairing possibilities. We used RNA bind-n-seq with purified AGO2–miRNA complexes to measure relative affinities of >1,000 3′-pairing architectures for each miRNA. In some cases, optimal 3′ pairing increased affinity by >500-fold. Some miRNAs had two high-affinity 3′-pairing modes—one of which included additional nucleotides bridging seed and 3′ pairing to enable high-affinity pairing to miRNA nucleotide 11. The affinity of binding and the position of optimal pairing both tracked with the occurrence of G or oligo(G/C) nucleotides within the miRNA. These and other results advance understanding of miRNA targeting, providing insight into how optimal 3′ pairing is determined for each miRNA.

**HIGHLIGHTS:**

- RNA bind-n-seq reveals relative affinities of >1,000 3′-pairing architectures
- Two distinct 3′-binding modes can enhance affinity, by >500-fold in some instances
- G and oligo(G/C) residues help define the miRNA 3′ segment most critical for pairing
- Seed mismatch identity can influence the contribution of compensatory 3′ pairing

## INTRODUCTION

MicroRNAs (miRNAs) are ∼22-nt regulatory RNAs that are processed from hairpin precursors. Upon processing, miRNAs associate with an Argonaute (AGO) protein and base-pair to sites within mRNAs to direct the destabilization and/or translational repression of these mRNA targets (Bartel, 2018; Jonas and Izaurralde, 2015). For most sites that confer repression in mammalian cells, pairing to miRNA nucleotides 2–7, referred to as the miRNA seed, is critical for target recognition, with an additional pair to miRNA position 8 or an A across from miRNA position 1 often enhancing targeting efficacy (Bartel, 2009; Lewis et al., 2005). Such sites with a perfect 6– 8-nucleotide (nt) match to the miRNA seed region (Figure 1A, left) are heuristically predictive of repression, with longer sites being more effective than shorter ones and more sites being more effective than fewer sites (Agarwal et al., 2015; Grimson et al., 2007). In addition, contextual features extrinsic to a site itself can influence targeting efficacy (Agarwal et al., 2015; Ameres et al., 2007; Brown et al., 2005; Grimson et al., 2007; Kedde et al., 2007, 2010; McGeary et al., 2019; Nielsen et al., 2007; Sætrom et al., 2007; Tafer et al., 2008; Wan et al., 2014).

**Figure 1.**
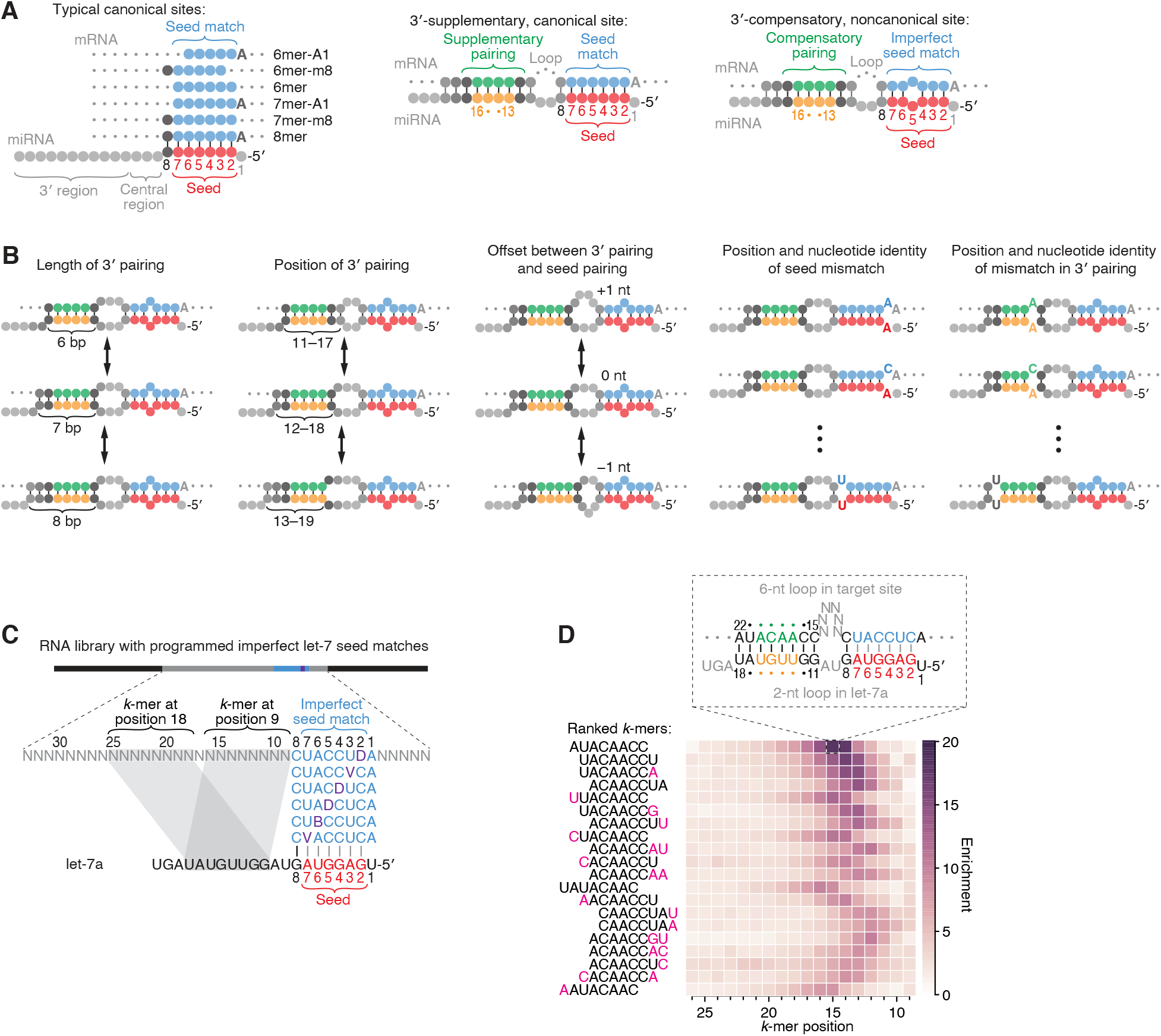
Features of miRNA 3′-compensatory sites characterized using AGO-RBNS. (**A**) Pairing of typical canonical sites (left), 3′-supplementary, canonical sites (middle), and 3′-compensatory, noncanonical sites (right). Canonical sites contain contiguous complementarity (blue) to the seed (red). Sites with shifted complementarity (i.e., the 6mer-A1 and 6mer-m8 sites) are sometimes also classified as canonical sites. 3′-supplementary sites have pairing to the miRNA 3′ region, which supplements canonical seed pairing and is reported to be most effective if it centers on miRNA nucleotides 13–16 (green and orange). This 3′ pairing can supplement 8mer sites (as shown) as well as other canonical sites (not shown). 3′-compensatory sites resemble 3′-supplementary sites, except they lack perfect pairing to the seed and thus pairing to the 3′ region helps to compensate for this imperfect seed pairing. Vertical lines represent Watson–Crick pairing. (**B**) The architectures of 3′ sites. Three independent features define each architecture: 1) the length of 3′ pairing (left), measured as the number of contiguous base pairs to the miRNA 3′ region; 2) the position of 3′ pairing (middle-left), defined as the 5′-most miRNA nucleotide engaged in 3′ pairing; and 3) the offset between the seed pairing and 3′ pairing (middle), which specifies the number of unpaired nucleotides separating the seed- and 3′-paired segments in the target RNA relative to that in the miRNA. Mismatches to the seed pairing (middle-right) or within the 3′ pairing (right) can elaborate on these architectures, as can bulged nucleotides (not shown). (**C**) A programmed RNA library for using AGO-RBNS to examine 3′ pairing of let-7a. The library contains an 8-nt region with all 18 possible single-nucleotide mismatches (purple) to the let-7a seed (red), with 25 nt of random-sequence RNA upstream of this region and 5 nt of random-sequence RNA downstream. *k*-mer positions are numbered with respect to the programmed 8-nt mismatched site. B represents C, G, or U; D represents A, G, or U; V represents A, C, or G; N represents A, C, G, or U. The black vertical line depicts perfect pairing at position 8, and gray vertical lines indicate Watson–Crick matches at only five of the six seed positions. (**D**) The top 20 8-nt *k*-mers identified by AGO-RBNS performed with the highest concentration of AGO2–let-7a (840 pM) and the programmed library (100 nM). *k*-mers were ranked by the sum of their enrichments at the five positions of the library at which they were most enriched. Left, alignment of *k-*mers, indicating in pink nucleotides that were not Watson–Crick matches to the miRNA. Right, heat map showing *k-*mer enrichment at each position of the library, with pairing shown for the top 8-nt *k-*mer at the position of its greatest enrichment. Black vertical lines depict perfect Watson–Crick pairing, and gray vertical lines indicate Watson–Crick matches at only five of the six seed positions.

Pairing to the miRNA 3′ region, particularly pairing that includes miRNA nucleotides 13– 16, can supplement perfect seed pairing to enhance targeting efficacy beyond that of seed pairing alone, and extensive pairing to the 3′ region can compensate for imperfect seed pairing to enable consequential repression (Brennecke et al., 2005; Grimson et al., 2007; Lewis et al., 2005). These two bipartite site types are referred to as 3′-supplementary and 3′-compensatory sites, respectively (Figure 1A, middle and right). Although 3′-supplementary sites are less common than sites with only a seed match, comprising ∼5% of all conserved sites observed in mammals, thousands of sites with preferentially conserved 3′-supplementary pairing are present in human 3′ UTRs (Friedman et al., 2009). Conserved 3′-compensatory sites are even less common, comprising only ∼1.5% of all preferentially conserved sites observed in human 3′ UTRs (Friedman et al., 2009). Nonetheless, two instances of this relatively rare site type within the 3′ UTR of *lin-41* mediate the extreme morphological and developmental defects by which the *let-7* miRNA was discovered in *C. elegans* (Ecsedi et al., 2015; Pasquinelli et al., 2000; Reinhart et al., 2000). Moreover, the use of these 3′-compensatory sites rather than canonical sites for *lin-41* repression is consequential; site mutations that create perfect seed pairing while maintaining the 3′ pairing cause precocious repression of the mRNA by other members of the let-7 seed family expressed during earlier larval stages (Brancati and Großhans, 2018). These results support the notion that 3′-compensatory sites enable differential target specificity between miRNAs that share a common seed sequence but differ within their 3′ regions (Brennecke et al., 2005; Lewis et al., 2005).

Although global analyses of site conservation and efficacy provide compelling evidence that pairing to the miRNA 3′ region is also utilized in mammalian cells (Friedman et al., 2009), these approaches have limitations for evaluating which 3′-pairing architectures are most effective: because some sites reside in mRNAs that respond indirectly to miRNA perturbation, analysis of site efficacy requires averaging results from multiple sites, which obscures identification of particularly efficacious sites. Likewise, because some sites are conserved by chance, analysis of site conservation requires averaging results from multiple sites to achieve statistical significance, which obscures identification of sites with unusual pairing architectures. Moreover, because each miRNA has relatively few 3′-supplementary sites and even fewer 3′-compensatory sites in the transcriptome, data from multiple miRNAs must be aggregated to observe a reliable signal of either efficacy or conservation, which prevents identification of miRNA-specific pairing preferences. Indeed, even when aggregating multiple miRNA-perturbation (e.g., transfection) datasets, which enables efficacy of 3′-supplementary sites to be detected (Grimson et al., 2007), a signal for the efficacy of 3′-compensatory sites has not been reported, underscoring the challenge of using global efficacy studies to learn about these sites.

Understanding the contribution of 3′ pairing to miRNA targeting efficacy is further complicated by the vast number of 3′-pairing architectures that are possible for a single miRNA sequence. The pairing architecture of a 3′-compensatory site can be described by five characteristics: 1) the length of contiguous pairing between the site and the miRNA 3′ region, 2) the position of pairing to the miRNA 3′ region, as defined by the 5′-most miRNA nucleotide involved in 3′ pairing, 3) the difference between the number of unpaired target nucleotides and the number of unpaired miRNA nucleotides bridging the seed and 3′ pairing, hereafter referred to as the “3′-pairing offset,” 4) the nature of the imperfect pairing to the seed, and 5) the nature of any imperfections in the 3′ pairing (Figure 1B). When considering only sites with perfect 3′ pairing with lengths ranging from 4–11 base pairs (bp) at all possible 3′ positions, offsets ranging from − 4 to +16 nt, and seed pairing interrupted by one of 18 possible single mismatches (or wobbles) to the 6-nt seed, there are >16,000 possible variants to the site architecture. For any miRNA under consideration, most of these variants are not present in the transcriptome, which limits the ability of global analyses of conservation or repression efficacy to determine which architectures are more effective than others.

The observation that miRNA targeting efficacy observed in the cell is largely a function of the affinity between AGO–miRNA complexes and their sites (McGeary et al., 2019) indicates that contributions of 3′ pairing to affinities measured in vitro can provide insight into biological targeting efficacy. Early measurements showed that pairing to positions 13–16 of let-7a imparts only a 2-fold increase in binding affinity, which led to the view that 3′-supplemental pairing contributes only modestly to affinity (Wee et al., 2012). Further measurements revealed some differences between miRNAs, with the observation that pairing to positions 13–16 of miR-21 increases affinity by 11-fold (Salomon et al., 2015), and a striking effect of longer pairing, with the observations that 10 bp of 3′-supplementary pairing to miR-122 and 9 bp of 3′-supplementary pairing (including a terminal G:U wobble) to miR-27a increases affinity by 20- and >400-fold, respectively (Sheu-Gruttadauria et al., 2019a). Other measurements illustrate the influence of the length of the target segment bridging the seed and 3′ pairing, with binding affinity varying ∼10-fold as this length is varied over a range of 1–15 nt (Sheu-Gruttadauria et al., 2019b). Taken together, these reports demonstrate the potential for miRNA 3′ pairing to enable high-affinity binding, and also illustrate that the benefit of this pairing varies, depending on the miRNA sequence and 3′-pairing architecture. Understanding how these features together modulate the benefit of 3′ pairing will be possible only after acquiring many more measurements with multiple miRNA sequences.

Imaging-based, high-throughput single-molecule biochemistry has recently been applied to acquire affinity measurements for ∼23,000 sites for each of two miRNAs (let-7a and miR-21), including many sites with 3′ pairing (Becker et al., 2019). These measurements revealed that miR-21 relies more on 3′ pairing when binding to a fully complementary target than does let-7a, that homopolymeric insertions are the least disruptive to binding when inserted between nucleotides 8 and 11 within the context of fully complementary binding, and that mismatches near the miRNA 3′ terminus (after position 16) increase target slicing and decrease binding affinity. However, because the design of target libraries was based primarily on fully complementary RNA targets to which varying extents of mismatched, bulged, and deleted nucleotides were introduced, only a small minority of the possible 3′-pairing architectures were queried. A fuller understanding of the contribution of pairing to the miRNA 3′ region requires many more affinity measurements with target RNA sequences that vary with respect to their seed pairing, and the position, offset, and length of 3′ pairing.

RNA bind-n-seq (RBNS) enables unbiased, high-throughput assessment of binding sites embedded within a larger random-sequence context (Dominguez et al., 2018; Lambert et al., 2014). We recently adapted RBNS for the study of miRNA targeting, and we built an analysis pipeline enabling calculation of relative equilibrium dissociation constants (*K*_D_ values) for many thousands of different RNA *k*-mers ≤12 nt in length (McGeary et al., 2019). Here, we further adapted the AGO-RBNS protocol to enable examination of sites >12 nt in length, thereby enabling the high-throughput investigation of bipartite sites containing near-perfect seed pairing and 4–11 additional pairs to the miRNA 3′ region. We applied this modified protocol to the systematic interrogation of the contribution of 3′ pairing for three natural miRNA sequences and four synthetic derivatives.

## RESULTS

### RBNS measures affinities for many 3′-compensatory sites of let-7a

AGO-RBNS begins with a series of 4–6 binding reactions, each containing an RNA library at a fixed concentration and a purified AGO–miRNA complex at a variable concentration spanning a 100-fold range (McGeary et al., 2019). Each molecule of the RNA library has a central region of random-sequence nucleotides flanked by constant sequences on each side that enable preparation of sequencing libraries. Upon reaching binding equilibrium, each reaction is passed through a nitrocellulose membrane, which retains AGO–miRNA complexes and any library molecules that are bound to the complexes. These bound library molecules are isolated and subjected to high-throughput sequencing, along with the input RNA library. Binding of an individual *k*-mer can be detected as enrichment in the bound compared to input sequences, and relative *K*_D_ values can be estimated simultaneously for hundreds of thousands of different *k*-mers by fitting a biochemical model to *k*-mer fractional abundances from each of the bound libraries.

As originally implemented, AGO-RBNS cannot provide reliable information on sites with more than ∼5 supplementary/compensatory pairs because such sites, which involve >12 bp of total pairing (Figure 1A, middle and right), are too rare in the sequences obtained from the input RNA library to enable accurate calculation of enrichment values. To overcome this constraint for sites to let-7a, a miRNA with physiologically relevant 3′ pairing (Brancati and Großhans, 2018; Pasquinelli et al., 2000; Reinhart et al., 2000), we used a library that contained a programmed region of imperfect seed pairing to let-7a, with 25 and 5 nt of random-sequence RNA separating the programmed region from the 5′ and 3′ constant sequences, respectively (Figure 1C). In each library molecule, this programmed region of imperfect seed pairing contained a let-7a 8mer site with a mismatch at one of its six seed nucleotides, such that each library molecule had one of 18 possible single-nucleotide seed mismatches (including wobbles) in approximately equal proportion. With this programmed region of imperfect seed pairing, each library contained 3′-compensatory sites at an ∼250-fold greater frequency than expected for a fully randomized RNA library.

AGO-RBNS was performed using this programmed library and purified AGO2–let-7a. For our initial analysis, we calculated the enrichment of all 8-nt *k*-mers at each position between the programmed region and the 5′-constant region of the library, after first removing reads with any of the six canonical sites to let-7a. The enriched *k*-mers had substantial complementarity to the 3′ region of let-7a (Figure 1D). The most enriched was AUACAACC—the perfect Watson– Crick match to positions 11–18 of the let-7a miRNA (Figure 1D). This 8-nt 3′ site was most strongly enriched when starting at position 15 of the library, which suggested that an internal loop with two miRNA nucleotides (9 and 10) and six target-site nucleotides (positions 9–14) separating seed pairing and 3′ pairing was optimal (Figure 1D, top). Using our nomenclature (Figure 1B), this 3′ site was classified as a position-11 site with pairing length of 8 bp and offset of +4 nt. This 8-nt, position-11 site was also ≥5-fold enriched at seven other neighboring offsets (corresponding to library positions 8–15), indicating that looping out 3–10 unpaired library nucleotides opposite miRNA nucleotides 9 and 10 was tolerated, albeit to varying degrees (Figure 1D).

The second-most enriched 8-nt *k*-mer was UACAACCU—the perfect Watson–Crick match to let-7a positions 10–17 (Figure 1D). This 3′ site had a maximal enrichment with five, rather than six, unpaired library nucleotides spanning the seed and 3′ pairing, with the distribution of enrichments shifted by 1 nt in comparison to that of the AUACAACC site. This 1-nt shift in the enrichment distribution corresponded with the 1-nt shift in site position (from 11 to 10 of the miRNA) to maintain an optimal offset of +4 target nucleotides. Indeed, the next 18 most enriched 8-nt *k*-mers represented 3′ sites with the pairing positions ranging from miRNA nucleotides 9–12, with enrichment distributions that correspondingly shifted to reflect an overall optimal offset of +4 target nucleotides (Figure 1D). Each had a contiguous stretch of 6–8 perfect Watson–Crick pairs to the let-7a 3′ region, usually including the ACAACC *k*-mer, which suggested that perfect pairing to let-7a positions 11–16, with a +4-nt offset, was particularly important for enhancing site affinity.

### let-7a has two distinct 3′-pairing modes

For a more comprehensive examination of 3′ sites of varied lengths, positions, and offsets (Figure 1B), we enumerated 3′ sites of lengths 4–11 nt that perfectly paired to the miRNA starting at any position downstream of nucleotide 8. For each length and position of 3′ pairing (e.g., for the 8mer-m11–18), we further enumerated all pairing offsets compatible with the 3′ site residing within the 25-nt random-sequence region upstream of the programmed site, converting each library position to an offset value based on the pairing position of each 3′ site (Figure 2A). For our initial *K*_D_ estimation and analyses, we pooled the reads for the 18 possible seed-mismatch types. This pooling increased the read counts for each 3′-pairing architecture, which enabled examination of sites as long as 11 nt, which in turn enabled analysis of 1006 distinct 3′-pairing architectures. We also enumerated and analyzed each canonical site (including the 6mer-m8 and 6mer-A1 sites, Figure 1A) residing within the 25-nt random-sequence region, as well as each of the 18 single-nucleotide seed-mismatch sites residing within this region, for use in benchmarking our results for 3′-compensatory sites.

**Figure 2.**
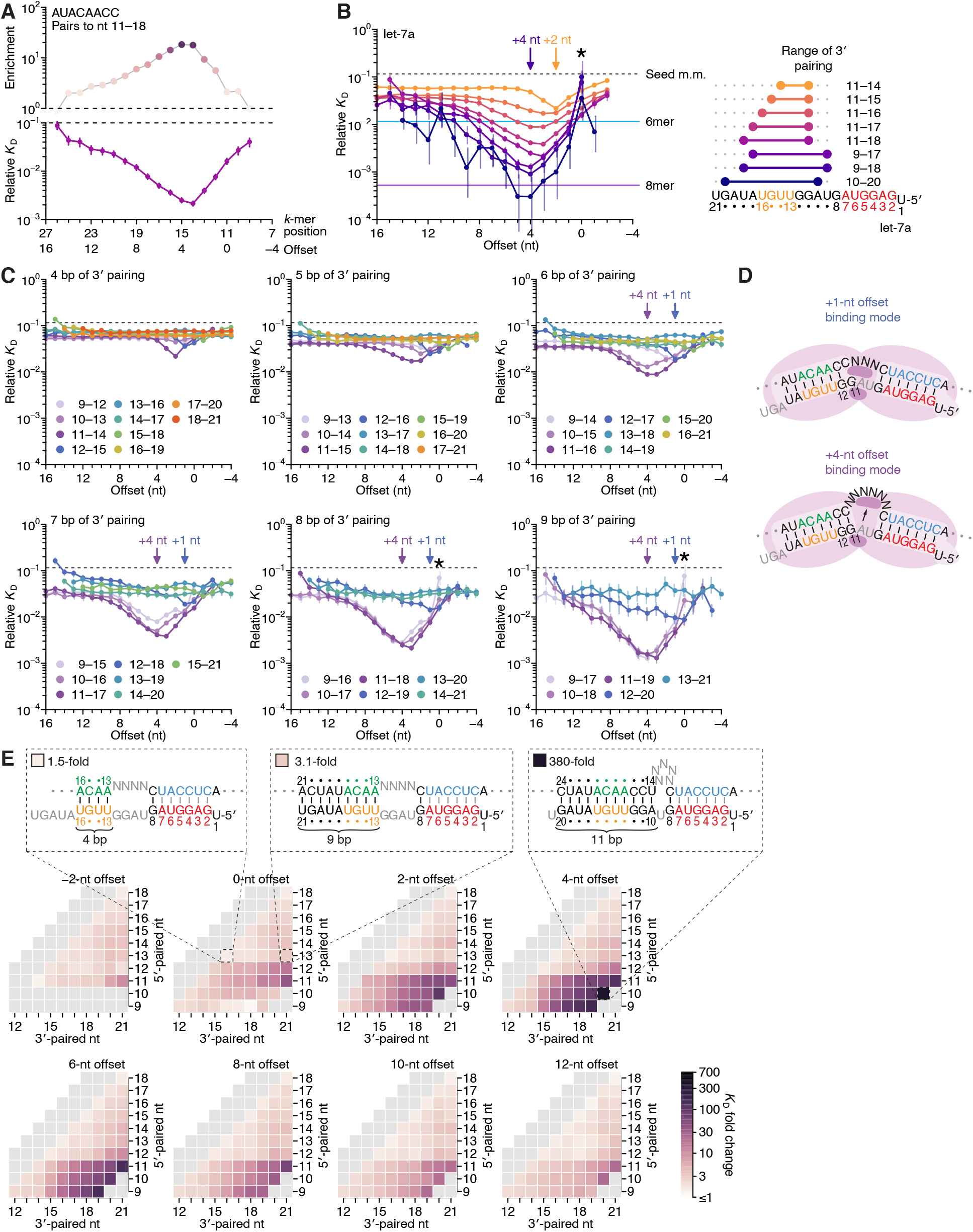
Pairing to nucleotide 11 and a +4-nt offset promote high-affinity binding to let-7a. (**A**) Correspondence of enrichment and relative *K*_D_ value of sites with the AUACAACC *k*-mer (the perfect match to miRNA positions 11–18) measured at each position in the programmed library. Each of these positions (upper *x*-axis) correspond to the indicated offset (lower *x*-axis). For example, because this *k*-mer paired to miRNA positions 11–18, pairing beginning at *k*-mer position 11 had a 0-nt offset. The *k*-mer enrichments and their associated colors (top) correspond to those of the top row of Figure 1D. (**B**) Relative *K*_D_ values of let-7a 3′-compensatory sites that had optimally positioned 3′ pairing of 4 (orange) to 11 (dark blue) bp. For each length of 3′ pairing, the optimal position is shown in terms of its complementarity to let-7a (right). For each of the 3′-compensatory sites, the relative *K*_D_ value is plotted as a function of its offset (left), as done previously for sites with 8 bp of optimally positioned 3′ pairing (Figure 2A). Vertical lines indicate 95% confidence intervals. The dashed horizontal line indicates the geometric mean of the 18 relative *K*_D_ values of the seed mismatch sites, each calculated from reads with <4 nt of contiguous complementarity to the miRNA 3′ region. The horizontal blue and purple lines indicate the relative *K*_D_ values of the canonical 6mer and 8mer sites, respectively. The arrows at +2 and +4 nt mark a shift in the optimal offset observed with increasing 3′ pairing length. The asterisk denotes the anomalously low binding affinity measured for pairing at position 9 with an offset of 0 nt. (**C**) The dependency of let-7a 3′-pairing affinity on pairing length, position, and offset. Each panel shows the relative *K*_D_ values of 3′-compensatory sites with 3′ pairing of a specified length over a range of positions and offsets. Each trend line is colored according to pairing position, spanning positions 9 (light violet) to 18 (red) when possible. The arrows at +1 and +4 nt mark a shift in the optimal offset as the position of 3′ pairing shifted to include nucleotide 11 of let-7a. Otherwise, these panels are as in (B, left). (**D**) Schematics of the two 3′-binding modes. In the +1-nt offset binding mode (top), miRNA nucleotide 11 is inaccessible due to occlusion by the central region of the AGO protein. In the +4-nt offset binding mode (bottom), the longer stretch of bridging target nucleotides enables a conformation in which nucleotide 11 is available for pairing to the target RNA. Although not intended to accurately reflect the conformation of either binding mode, these schematics illustrate how a larger offset might enable pairing to a more centrally located miRNA nucleotide. (**E**) Affinity profile of the let-7a 3′ region. Each cell indicates the fold change in relative *K*_D_ attributed to a 3′ site with indicated length, position, and offset of pairing. Each row within a heat map corresponds to a different miRNA nucleotide at the start of the 3′ pairing, and each column corresponds to a different miRNA nucleotide at the end of the 3′ pairing. Each heat map shows the results for a different offset. The three diagrams indicate the fold-change values and architectures for 3′ sites pairing to miRNA nucleotides 13–16 with an offset of 0 nt (left), pairing to miRNA nucleotides 13–21 with an offset of 0 nt (middle), and pairing to miRNA nucleotides 10–20 with an offset of +4 nt (right). Gray boxes indicate pairing ranges that were either too short (<4 bp) or too long (>11 bp) for relative *K*_D_ values to be reliably calculated. Black vertical lines depict perfect Watson–Crick pairing, and gray vertical lines indicate Watson–Crick matches at only five of the six seed positions.

Simultaneous estimation of the fractional abundance of these sites in each of the AGO2–let-7a-bound libraries in comparison to that of the input library enabled calculation of their relative *K*_D_ values. As illustrated for the 8-nt *k*-mer identified as most enriched in the previous analysis (Figure 1D, top row), variation in *K*_D_ values qualitatively tracked with that of enrichment values but quantitatively differed due to the attenuating effects of background binding and site saturation on enrichment values (McGeary et al., 2019) (Figure 2A). Relative *K*_D_ values corresponding to a broad spectrum of 3′ pairing architectures spanned a >500-fold range, with strong agreement observed between the results of replicate experiments performed independently with different preparations of both AGO2–let-7a and the let-7a programmed library (*r*^2^ = 0.96, *n* = 1477; Figure S1A, left). Agreement between the two replicates was maintained, albeit to a lesser degree, when read counts for each 3′-pairing architecture were not pooled over the 18 seed-mismatched sites in the programmed region (*r*^2^ = 0.78, *n* = 23,912; Figure S1A, right). Furthermore, for shorter 3′ sites, which could be analyzed using data from a standard AGO-RBNS experiment that used a non-programmed random-sequence library (McGeary et al., 2019), the relative *K*_D_ values determined from the programmed library correlated well with those determined from a random-sequence library (*r*^2^ = 0.83, Figure S1B). Despite the overall correlation, a minor systematic difference in the values for the same sites determined from the two types of libraries was observed. This distortion was presumed to be due to the absence of no-site-containing RNA molecules in the programmed library and was corrected accordingly (Figure S1B).

To investigate the interplay of pairing position, length, and offset, we identified the optimal 3′ sites of lengths 4–11 nt and, as in Figure 2A, examined the effect of varying offset on the affinity of each of these sites (Figure 2B). Nearly all possibilities examined had values readily distinguished from the log-averaged value for seed-mismatched sites alone, with compensatory pairing to miRNA nucleotides 11–16 at optimal offsets yielding binding affinities comparable to that of the canonical 6mer (Figure 2B, left). Further inspection of longer 3′ sites underscored the conclusion that pairing to the GGUUGU segment spanning positions 11–16 of let-7a is the most consequential for 3′-compensatory pairing, as all optimal pairing positions for 3′ sites ≥6 nt in length paired to this segment. Moreover, inspection of the optimal positions for shorter sites showed that pairing to the 5′ end of this segment (containing the sequence GGUU) was more impactful than pairing to its 3′ end (Figure 2B, right). In addition, increasing the length of pairing from 4 to 11 bp led not only to increased binding affinity at almost all offsets, as might have been expected, but also a shift in the optimal offset, with a preferred offset of +2 nt when pairing with 4 bp compared to a preferred offset of +4 nt when pairing with 9–11 bp (Figure 2B, left).

To investigate the underpinnings of the change in preferred offset, we plotted the relative affinities of all possible positions, lengths, and offsets for let-7a 3′ pairing (Figure 2C). As pairing length increased beyond 6 bp, two distinct trends emerged: one with a preferred offset of +4 nt (or sometimes +3 nt) and higher-affinity relative *K*_D_ values, and another with a preferred offset of +1 nt and more modest relative *K*_D_ values. These two offset trends indicated two distinct binding modes. Moreover, the preferred offset of +4 nt nearly always occurred for configurations that included pairing to the G at position 11 of let-7a, with a switch from the preferred offset of +1 nt to a preferred offset of +4 nt when 3′ pairing began at position 11 rather than 12. These results suggested that pairing to position 11 in the central region of the miRNA is less accessible than pairing to position 12, and therefore a longer loop in the target sequence is required to bridge seed pairing with 3′ pairing that includes position 11 (Figure 2D). Nonetheless, when the increased offset enables pairing to position 11 through this second binding mode, substantially greater affinity can be achieved.

Some of the lowest relative affinity values were observed for extended 3′-pairing possibilities that began at position 9 with an offset of 0 nt (Figures 2B and 2C, asterisks). These low values were attributed to AGO2-catalzyed slicing of molecules with extensive contiguous pairing, which depleted these molecules from our bound library. Supporting this idea, analogous sites with offsets of either − 1 or +1 nt, which were expected to disrupt slicing due to single-nucleotide bulges in either the miRNA or the site, respectively, did not have aberrantly low relative affinities. Our observation of some slicing during the course of the binding experiments agreed with reports that AGO2 can slice sites that have a seed mismatch but are otherwise extensively paired to the guide RNA (Becker et al., 2019; Chen et al., 2017; Wee et al., 2012).

We next used heat maps to visualize the interplay between 3′-site position and pairing length at different offsets (Figure 2E). Within each heat map, adjacent cells corresponded to the difference in *K*_D_ fold change caused by the addition or removal of a pair at either the 5′ end (adjacent rows) or the 3′ end (adjacent columns) of the 3′ site, while maintaining the same offset. For example, in the heatmaps corresponding to offsets of +4 to +12 nt, the prominent contrast between the row corresponding to pairing beginning with nucleotide 11 and the row corresponding to pairing beginning with nucleotide 12 illustrated the strong benefit of pairing to G11 of let-7a (Figure 2E). At the optimal offset length of +4 nt, pairing to let-7a positions 10–20 conferred an ∼380-fold increase in affinity over the average seed-mismatched site alone (Figure 2E), leading to an overall binding affinity rivaling that of the canonical 8mer (Figure 2B). The binding affinity of this site and all other sites decreased nearly uniformly as offset values increased beyond +4 nt. Binding affinity decreases were less uniform as offset values decreased to 0 and –2 nt, which reflected a switch from the +4-nt binding mode to the +1-nt binding mode, with a concomitant reduction in the benefit of pairing to nucleotide 11.

Previous low-throughput measurements of the benefit of 3′ pairing for let-7a examined the influence of pairing to miRNA positions 13–16 at an offset of 0 nt and found that this pairing conferred a 1.6–2-fold increase in binding affinity (Salomon et al., 2015; Wee et al., 2012). Likewise, our measurements for this 4-nt 3′ site indicated that it conferred a 1.5-fold increase in affinity (Figure 2E). Furthermore, maintaining the offset of 0 nt and the pairing position of 13 and extending pairing to the very 3′ end of let-7a improved the binding affinity to only 3.1-fold (Figure 2E). These results highlight the importance of both the +4-nt offset and pairing to position 11 of let-7a—two features that would have been difficult to identify without comprehensive investigation of the 3′-pairing preferences of this miRNA. Indeed, the importance of these two features is not revealed in an analysis of a dataset that reports the affinities of ∼23,000 different sites to let-7a, because these ∼23,000 sites were not designed to analyze the combined effects of varying both pairing position and pairing offset (Becker et al., 2019) (Figure S2).

### Different miRNAs have distinct 3′-pairing preferences

The optimal 3′-pairing architecture for let-7a differed from that previously elucidated for miRNAs more generally (Grimson et al., 2007). When pooling repression and conservation data for 11 miRNAs, pairing to miRNA nucleotides 13–16, with an offset of 0 nt appears to be most consequential (Figure 1A) (Grimson et al., 2007). Because the previous analysis represents the average of trends derived from multiple miRNAs, a diversity of miRNA-specific 3′-pairing preferences might explain this disagreement. We therefore measured the 3′-pairing profiles of two other well-studied miRNAs, miR-1 and miR-155, for comparison to the let-7a profile.

Stabilizing 3′ pairing was observed for both miR-1 (Figures 3A) and miR-155 (Figures 3B), with binding affinity increasing with the length of pairing, as observed for let-7a (Figure 2). However, the magnitude of increased binding affinity differed from that of let-7a and that of each other: the affinity of 3′ pairing to miR-1 was more modest, with 3′-compensatory sites seldomly reaching the affinity of its canonical 6mer site (Figure 3A), whereas for miR-155, they often reached the affinity of its canonical 8mer site (Figure 3B). The positions of the best sites at each length also differed from let-7a. For miR-1, optimal 4-nt sites paired to miRNA nucleotides 12– 15, and when considering optimal sites of increasing lengths, pairing extended continuously, primarily towards the 3′ end of the miRNA and never reaching miRNA nucleotide 10 (Figure 3A, right). By contrast, for miR-155, optimal 4-nt sites paired to miRNA nucleotides 13–16, and for optimal sites of increasing lengths, pairing sometimes shifted discontinuously and never included miRNA nucleotide 12 (Figure 3B, right).

**Figure 3.**
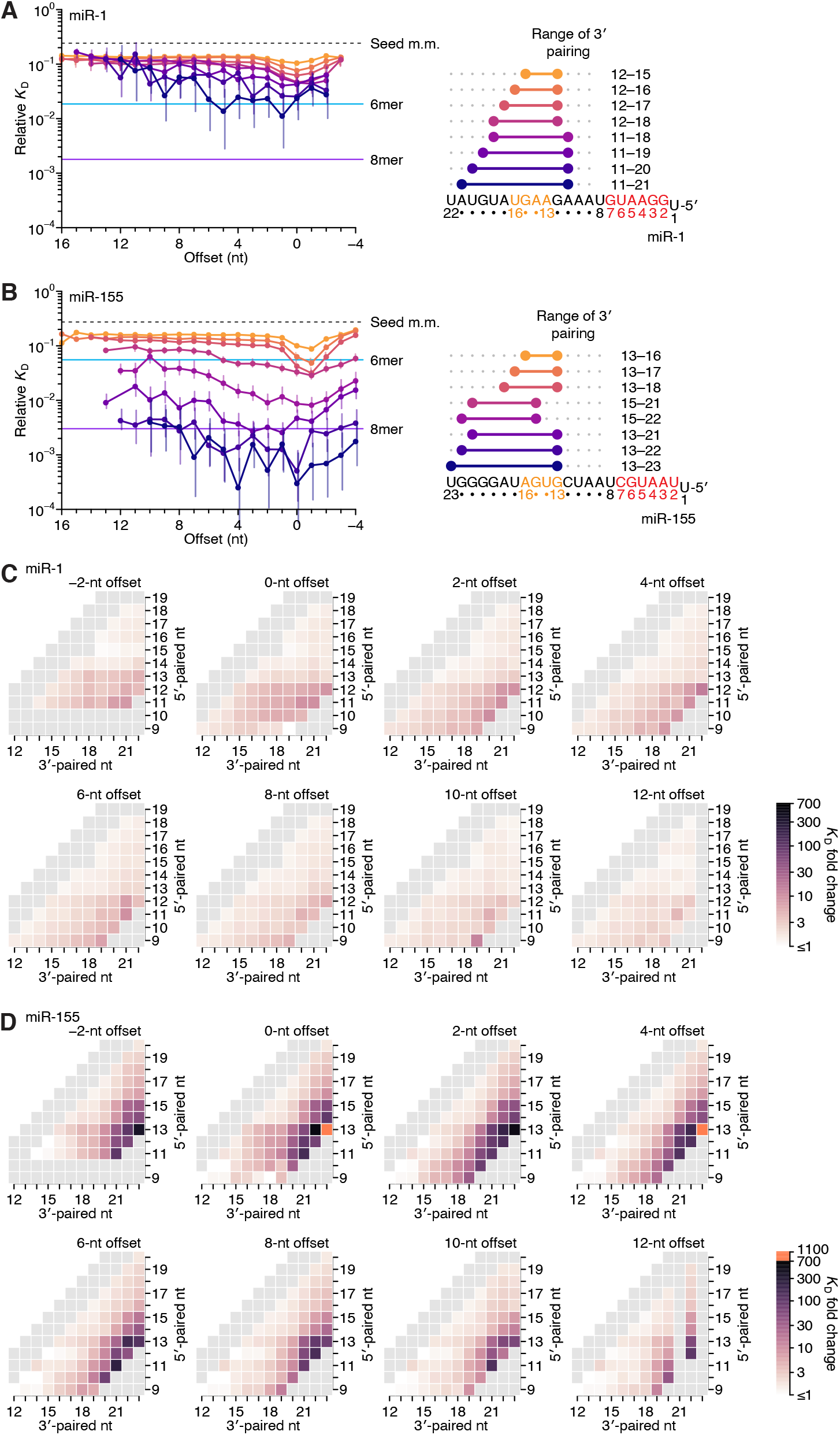
Relative affinity measurements of 3′-compensatory sites of miR-1 and miR-155. (**A**) Relative *K*_D_ values of miR-1 3′-compensatory sites that had optimally positioned 3′ pairing of 4–11 bp. Otherwise, this panel is as in Figure 2B. (**B**) Relative *K*_D_ values of miR-155 3′-compensatory sites that had optimally positioned 3′ pairing of 4–11 bp. Otherwise, this panel is as in Figure 2B. (**C** and **D**) Affinity profiles of the 3′ regions of miR-1 (C) and miR-155 (D). Otherwise, these panels are as in Figure 2E.

Analysis of each of the optimal 3′ sites of miR-1 and miR-155 along the length of the random region indicated that, unlike sites for let-7a, those for neither of these two miRNAs underwent a significant shift in the preferred offset (Figures 3A and 3C, left). Nevertheless, the longer optimal sites of miR-1 extended to position 11, and their range of near-optimal offsets broadened to include values from 0 to +5 nt, consistent with contributions from both binding modes. The offset preferences of miR-155 also broadened with increased pairing. However, instead of coinciding with pairing at position 11, these broadened preferences coincided with pairing to the G19-G20-G21-G22 stretch near the 3′ end of miR-155. These results might relate to the ability of this miRNA to participate in seed-independent 3′ pairing, as detected when performing AGO-RBNS with fully randomized RNA libraries (McGeary et al., 2019). However, our data were not suitable for studying seed-independent pairing, due to the presence of an imperfect 8mer site in each molecule of the programmed library.

In summary, the most optimal 3′ sites each paired to at least two nucleotides of the miRNA segment spanning positions 13–16, which was previously identified as most consequential for 3′ pairing, but frequently did not pair to the entire segment. Shorter optimal sites consistently preferred pairing to G nucleotides adjacent to miRNA positions 13–16. For example, shorter optimal sites to let-7a paired to the G11-G12 sequence element 5′ of this segment rather than to G15-U16 (Figure 2B, right), the optimal 4-nt site to miR-1 paired to G12 rather than to U16 (Figure 3A, right), and intermediate-length optimal sites to miR-155 paired to G19-G20-G21 rather than to G13-U14 (Figure 3B, right). These trends were also observed when examining many combinations of positions, lengths, and offsets for miR-1 and miR-155 (Figure S3). In aggregate, these results supported the report of an intrinsic preference for pairing to miRNA nucleotides 13–16 (Grimson et al., 2007) but also indicated that the miRNA sequence imparts additional preferences, resulting in unanticipated differences between the optimal sites of individual miRNAs. These sequence-specific preferences tended to favor pairing to G residues of the miRNA, which was presumably explained by the greater stability of G:C pairing over A:U pairing, although the presence of only a single C nucleotide prevented investigation of whether pairing to G was preferred over pairing to C. We also observed differences between miRNAs in the strength of 3′ pairing. Compared to 3′-site affinities observed for let-7a, affinities were substantially lower for miR-1 and substantially greater for miR-155 (median increase in affinity with 11 bp of 3′ pairing of 36-fold, 5.8-fold, and 133-fold for let-7a, miR-1 and miR-155, respectively). Thus, our results indicated that association of the guide RNA with the AGO protein does not fully standardize either the architecture of optimal 3′ pairing or the magnitude of its benefit.

### Pairing and offset coefficients describe unique 3′-pairing profiles for each miRNA

To summarize the results for miR-1 and miR-155, we generated heat maps representing the binding affinity at all possible pairing positions for all pairing lengths of 4–11 bp, as a function of pairing offset (Figures 3C and 3D), as with let-7a (Figure 2E). The similarities observed between heat maps for the same miRNA at different offsets indicated that each change in offset altered the binding affinity of all 3′-pairing possibilities in a consistent manner, which in turn indicated that for each of the three miRNAs, the effect of pairing offset was largely independent of the effect of guide–target complementarity (Figures 2E, 3C, and 3D). This overall independence was observed for let-7a, despite its two binding modes, because the contribution of the +4 binding mode, which had the higher affinities, dominated over that of the +1 binding mode.

To test this independence, we examined how well the affinities could be explained as the product of two coefficients, one representing the contribution of the pairing range, which was defined by pairing position and length (represented by the location of a cell within the heat maps of Figures 2E, 3C, and 3D), and the other representing the contribution of the pairing offset. Our model fit the data well (*r*^2^ = 0.92, 0.86, and 0.96 for let-7a, miR-1, and miR-155, respectively; Figure S4), and yielded a set of pairing and offset coefficients for each miRNA. Each pairing coefficient represented the Δ*G* of the corresponding pairing range at its optimal offset, and each offset coefficient represented the reduction in Δ*G* observed at suboptimal pairing offsets (Figures 4A–4C). For each miRNA, the pairing coefficients corresponded well with the affinities observed at the preferred offset (Figures 4A–4C, middle-right and right; *r*^2^ = 0.98, 0.97, and 0.96, respectively). Moreover, these coefficients, which distilled the pairing preferences indicated by the 934, 1061, and 1180 *K*_D_ fold-change values measured for let-7a, miR-1, and miR-155, respectively, quantitatively captured the qualitative observations made earlier from analysis of subsets of the data. For example, they captured the respective importance of pairing to nucleotides 11, 12, and 20 for let-7a, miR-1, and miR-155, as well as the respective preferences for offsets of +4, +1, and +1 nt. They also captured the more narrowed offset preferences of let-7a in comparison to those of miR-1 and miR-155 (Figures 4A–4C, middle-left) and the contribution of pairing starting at miRNA position 15 for miR-155 (Figure 4C, left).

**Figure 4.**
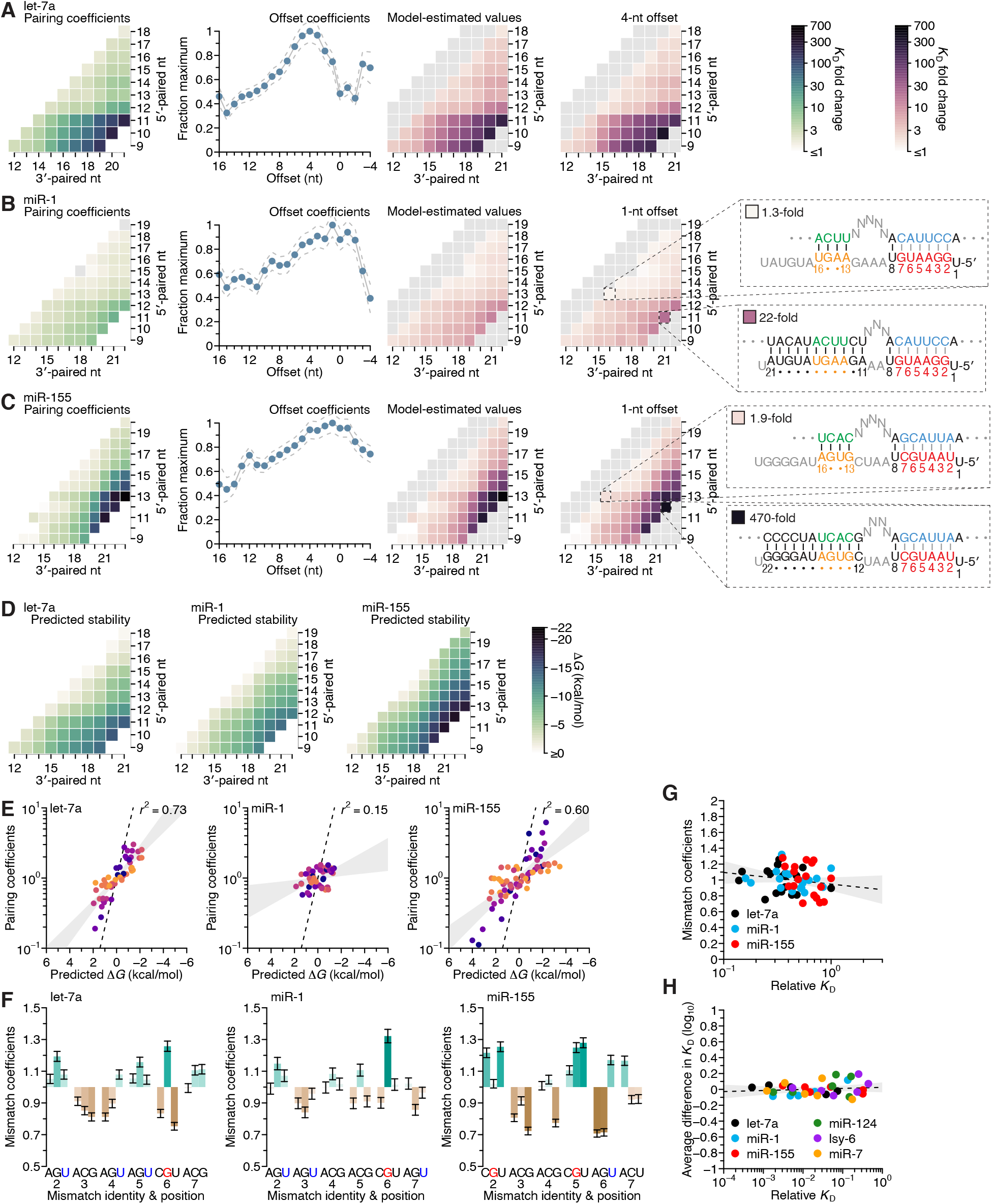
Distinct pairing, offset, and seed-mismatch preferences of different miRNAs. (**A**–**C**) Model-based analyses of 3′-pairing preferences of let-7a (A), miR-1 (B), and miR-155 (C). For each miRNA, 3′-pairing affinities are described by a set of pairing coefficients (left) and offset coefficients (middle-left; dashed lines, 95% confidence interval), which when multiplied together (middle-right) approximated measured *K*_D_ fold-change values (right; let-7a values replotted from Figure 2E). The parameters were obtained by maximum-likelihood estimation with a nonlinear energy model. For both miR-1 (B) and miR-155 (C), the two pairing diagrams indicate the fold-change value and architecture for a 3′ site pairing to miRNA nucleotides 13–16 (top) in comparison to the fold-change value and architecture of the 3′ site with the greatest measured affinity (bottom) at their shared optimal offset of +1 nt. Pairing coefficients, model predictions, and *K*_D_ fold-change values of miR-1 were not calculated for pairing to miRNA positions 15–18 and 19–22 because these two segments were identical (gray boxes). (**D**) Predicted Δ*G* values of the 3′ sites with pairing ranges in (A–C). (**E**) The relationship between the model-derived pairing coefficients (A–C) and the predicted Δ*G* values (D). Points are colored according to pairing length, as in Figure 2A. To control for the trivial effect of increasing pairing length, pairing coefficients were divided by the geometric mean of all coefficients with the same length, and Δ*G* values of each length were normalized to the mean Δ*G* value of pairings with the same length. The gray region represents the 95% confidence interval of the relationship when fitting a linear model to the data (*r*^2^, coefficient of determination), and the dashed line represents the predicted thermodynamic relationship given by *K* = e^− Δ*G*/*RT*^. (**F**) Distinct effects of seed-mismatches on 3′-pairing affinities of let-7a, miR-1, and miR-155. For each miRNA, seed-mismatch coefficients were derived by maximum-likelihood estimation, fitting a nonlinear model to the *K*_D_ fold-change values observed when examining 3′-site enrichment separately for each of the 18 seed mismatches. The error bars indicate 95% confidence intervals. Wobble pairing in which the G was in either the miRNA or the target is indicated in blue and red, respectively. (**G**) Relationship between affinity of 3′-compensatory pairing and that of seed-site binding. For each seed mismatch, the coefficient from (F) is plotted as a function of the relative *K*_D_ value of that mismatch, as measured using results from the programmed libraries for let-7a (black), miR-1 (blue), and miR-155 (red). The dashed line shows the linear least-squares fit to the data, with the gray interval indicating the 95% confidence interval. (**H**) Relationship between affinity of 3′-supplementary pairing and that of seed-site binding. For each of the six seed-matched site types (Figure 1A, left) and for each of the six miRNAs (key), the relative affinity of the top quartile of all 4- and 5-nt 3′ sites with their preferred offsets is plotted as a function of the relative affinity of the seed-matched site. Relative affinities were measured from analysis of previous AGO-RBNS that used a random-sequence library (McGeary et al., 2019) (Figure S11).

Because the pairing coefficients represented the thermodynamic benefit of each pairing possibility, we examined how well each set of pairing coefficients was explained by the nearest-neighbor model that predicts the stability of RNA hybridization in solution. To do so, we calculated the predicted Δ*G* value for each 3′ site (Figure 4D) and adjusted each value by subtracting the mean value for that length of pairing, which was done to remove the trivial effect of increasing pairing length (Figure 4E). When comparing these length-adjusted values with analogously adjusted pairing coefficients, we observed a strong relationship for both let-7a and miR-155, which explained most of the variation in the length-adjusted coefficients, and a much weaker relationship for miR-1. Nevertheless, even when focusing on results for let-7a and miR-155, the apparent effect size was less than that expected by the relationship Δ*G* = − *RT* ln*K* (Figure 4E, dashed lines). Thus, as observed with the miRNA seed region (McGeary et al., 2019; Salomon et al., 2015), compared to RNA free in solution, association with AGO reduces the differences in binding energy observed when hybridizing to different miRNA 3′-end sequences.

This reduction in magnitude also applied to the overall contribution of 3′ pairing (Figure S5A). For instance, although the >200-fold differences in binding affinity imparted by the top 11-nt 3′ sites of let-7a and miR-155 might seem large, the Δ*G* predicted for each of these sites was − 14.8 kcal/mol and − 20.1 kcal/mol, which corresponded to respective fold differences in affinity of 2.7 × 10^10^ and 1.5 × 10^14^. Presumably the benefit of pairing to 3′ sites was mostly offset by the cost of disrupting favorable interactions between unpaired 3′ regions and AGO, as proposed in the context of fully paired sites (Tomari and Zamore, 2005). The magnitude of this inferred cost appeared specific to each miRNA, implying that AGO might have some sequence preferences when interacting with unpaired miRNA 3′ regions. For example, pairing to either nucleotides 9–19 of let-7a or nucleotides 11–21 of miR-1 was predicted to occur with equivalent Δ*G* values of − 13.5 kcal/mol, yet the model-determined contributions of these sites were 160- and 14-fold, respectively (Figure S5A, left and middle).

Separating the comparison between *K*_D_ fold-change and Δ*G* based on whether the contiguous range of pairing included the G11, G12, and G20 of let-7a, miR-1, and miR-155, respectively, revealed a cooperative benefit of pairing to these nucleotides (Figures S5B and S5C), such that their inclusion within the 3′ pairing enabled the other paired nucleotides to contribute more to the interaction. We also note that using the measured affinities rather than pairing coefficients did not increase agreement with Δ*G* (Figures S5D and S5E), suggesting that the use of the pairing coefficients did not lead to loss of information contained within the data from which they were generated.

Having acquired these 3′-pairing profiles from the programmed libraries, we examined the extent to which such profiles could be generated using data obtained previously from fully randomized libraries (McGeary et al., 2019) (Figures S6A–S6F). For let-7a, miR-1, and miR-155, the pairing and offset coefficients derived from data from the two types of libraries agreed well with each other, provided that 3′-pairing lengths did not extend beyond 8 bp (Figures S6G and S6H). However, when pairing lengths extended beyond 8 bp, affinity values were not reliably determined because the sites were only sparsely represented in the random-sequence libraries.

We next turned to the analyses of miR-124, lsy-6, and miR-7, for which data from programmed libraries were not available. miR-124, like let-7a, had both preferred pairing to position 11 and an optimal pairing offset of >2 nt (Figure S6D). To look for evidence of multiple binding modes, we repeated the analyses of both Figure 2B (for pairing lengths of 4–8 bp) and Figure 2C (for pairing lengths of 4 and 5 bp), using the prior AGO-RBNS data for miR-124, lsy-6, and miR-7 (Figure S7). For comparison, we also repeated these analyses using the prior AGO-RBNS data for let-7a, for which we had evidence of two binding modes from the programmed-library AGO-RBNS data. For each of the four miRNAs, we found evidence of two binding modes. Both let-7a and miR-124 had the previously observed pattern, in which the binding mode with the positive offset and pairing to nucleotide 11 had binding affinity greater than that of the binding mode with an offset of 0-nt and pairing to only nucleotide 12 (Figures S7A–S7D). However, lsy-6 and miR-7 had a different pattern, in which the binding mode corresponding to the positive offset and pairing to nucleotide 11 had binding affinity similar to that of the binding mode with an offset of 0 nt and pairing to only nucleotide 12 (Figures S7E– S7H).

These examples further supported the proposal of a second binding mode, in which productive 3′ pairing extends to nucleotide 11, provided that additional unpaired target nucleotides are available to bridge pairing between the seed and this nucleotide. These results also suggested that pairing to the G11-G12 dinucleotide found in both the let-7a and miR-124 sequences enabled this second binding mode to dominate over the first, whereas pairing to the single G11 found in lsy-6 and miR-7 added to site affinity but did not enable the second binding mode to dominate. Indeed, although miR-7 appeared to have both binding modes, it had the weakest 3′-compensatory pairing of the six miRNAs profiled, with 8-nt 3′ sites never contributing more than an 18-fold increase in binding affinity.

The analyses of the miR-124 and lsy-6, which each had multiple C nucleotides in their 3′ region, allowed us to return to the question of whether pairing to miRNA G nucleotides might be favored over pairing to C nucleotides. Pairing to C15 of lsy-6 substantially added to binding affinity. For example, the 4.2-fold greater affinity of the position 12–15 site over the position 11– 14 site indicated that pairing to C15 was favored over pairing to G11, and extending pairing from positions 11–14 to 11–15 increased affinity 8.2-fold (Figure S6E). Pairing to C13 was also somewhat preferred, as illustrated by the 1.8-fold greater affinity of the 13–17 site over the 14– 18 site, and the 3.2-fold benefit of extending pairing from positions 14–18 to 13–18. However, pairing to C19-C20 of miR-124 did not seem to have the same impact as pairing to G19-G20 of miR-155, as illustrated by the negligible (0.9-fold) benefit of extending the miR-124 pairing from positions 13–18 to 13–20, compared to the 14-fold benefit for miR-155. These results supported the idea that pairing to a G in the miRNA 3′ region is generally favored over pairing to a C, although pairing to a C located centrally within the 3′ region can be impactful.

### The type of seed mismatch affects the affinity of 3′ pairing

To examine the influence of seed-mismatch position and identity, we analyzed the full set of 16,235, 18,076, and 19,666 relative *K*_D_ values of let-7a, miR-1, and miR-155, no longer combining read counts for the 18 possible seed-mismatch sites in the programmed library prior to *K*_D_ estimation. For each pairing, offset, and seed-mismatch possibility, the relative *K*_D_ value of the 3′-compensatory site was divided by that of its seed-mismatch site to generate a fold-change value representing the contribution of the 3′ site to affinity (Figures S8–S10). These values revealed a striking effect of seed-mismatch identity on the benefit of 3′ pairing. This effect was of greater magnitude for more favorable 3′ sites, causing affinities to vary >10-fold for the most optimal sites to miR-155. To further study this effect, we expanded our model to include a seed-mismatch coefficient, such that each log_10_(*K*_D_ fold change) value was described as the product of the pairing, offset, and seed-mismatch coefficients corresponding to its 3′-pairing architecture (Figures S8–S10).

The affinity of seed-mismatch sites lacking 3′ pairing had little relationship with the influence of the mismatch on 3′-pairing affinity (Figure 4G). Likewise, examination of data from the six random-library AGO-RBNS experiments found no relationship between the affinities of canonical sites lacking 3′ pairing and the relative influence of each canonical site on the benefit of supplemental pairing (Figures 4H and S11). Furthermore, the average effect of canonical-site type on 3′ binding affinity was small, with only six out of the 36 miRNA–site combinations having a >0.1 effect on log_10_(*K*_D_ fold change), corresponding to an ∼25% change in binding affinity (Figure 4H). Together, these results indicate that for 3′-supplementary pairing, the benefit of the 3′ pairing is largely the same between sites, but that for 3′-compensatory pairing, the potential benefit of 3′ pairing differs depending on the identity of the seed mismatch. This might be due to a differential ability of these mismatches to elicit a conformational change in AGO allowing pairing to the 3′ end (Schirle et al., 2014; Sheu-Gruttadauria et al., 2019b). Another potential contribution might stem from variation in elemental rate constants of seed-mismatch sites of similar affinity, whereby some sites have dwell times that are too short to establish pairing to the miRNA 3′ region.

When comparing the effects for guide–target nucleotide possibilities, strong trends did not emerge within miRNAs (e.g., when comparing the effects of mismatches to the G at position 2 with those of the mismatches to the G at position 4 of let-7a), or between miRNAs (e.g., when comparing the effects of mismatches to the G at position 3 of miR-1 with those to the G at position 6 of miR-155) (Figure 4F). However, in cases in which the same nucleotide occurred at the same position for two different miRNAs, some correspondence was observed (positions 2 and 6 of let-7a and miR-1, position 3 of let-7a and miR-155, position 4 of miR-1 and miR-155) (Figure 4F). Notably, the miRNA–target U:G mismatch at position 6, which was the most favored mismatch for both let-7 and miR-1, occurs within one of the two compensatory sites within the 3′ UTR of *C. elegans lin-41* (Pasquinelli et al., 2000; Reinhart et al., 2000).

### The seed-mismatch and 3′-sequence effects act independently

The distinct pairing, offset, and seed-mismatch preferences of the three miRNAs measured using the programmed libraries raised the question of the extent to which these preferences depended on the sequence of the seed region, the sequence of the 3′ region (i.e., beginning at miRNA nucleotide 9), or a combination of the two. To answer this question, we generated two chimeric miRNAs, one fusing the seed of let-7a to the 3′ region of miR-155 (let-7a–miR-155) and the other fusing the seed of miR-155 to the 3′ region of let-7a (miR-155–let-7a) (Figure 5A), and then performed AGO-RBNS using their corresponding seed-mismatched programmed libraries. As done previously with the natural miRNAs, we first determined the pairing and offset preferences of both chimeric miRNAs by summing over all 18 seed-mismatch types, measuring the *K*_D_ fold change for each range of pairing and offsets possible in the libraries, and fitting a multiplicative model of pairing and offset-preferences to the resultant 1181 and 934 measured affinity values for let-7a–miR-155 (Figure 5B) and miR-155–let-7a (Figure 5C).

**Figure 5.**
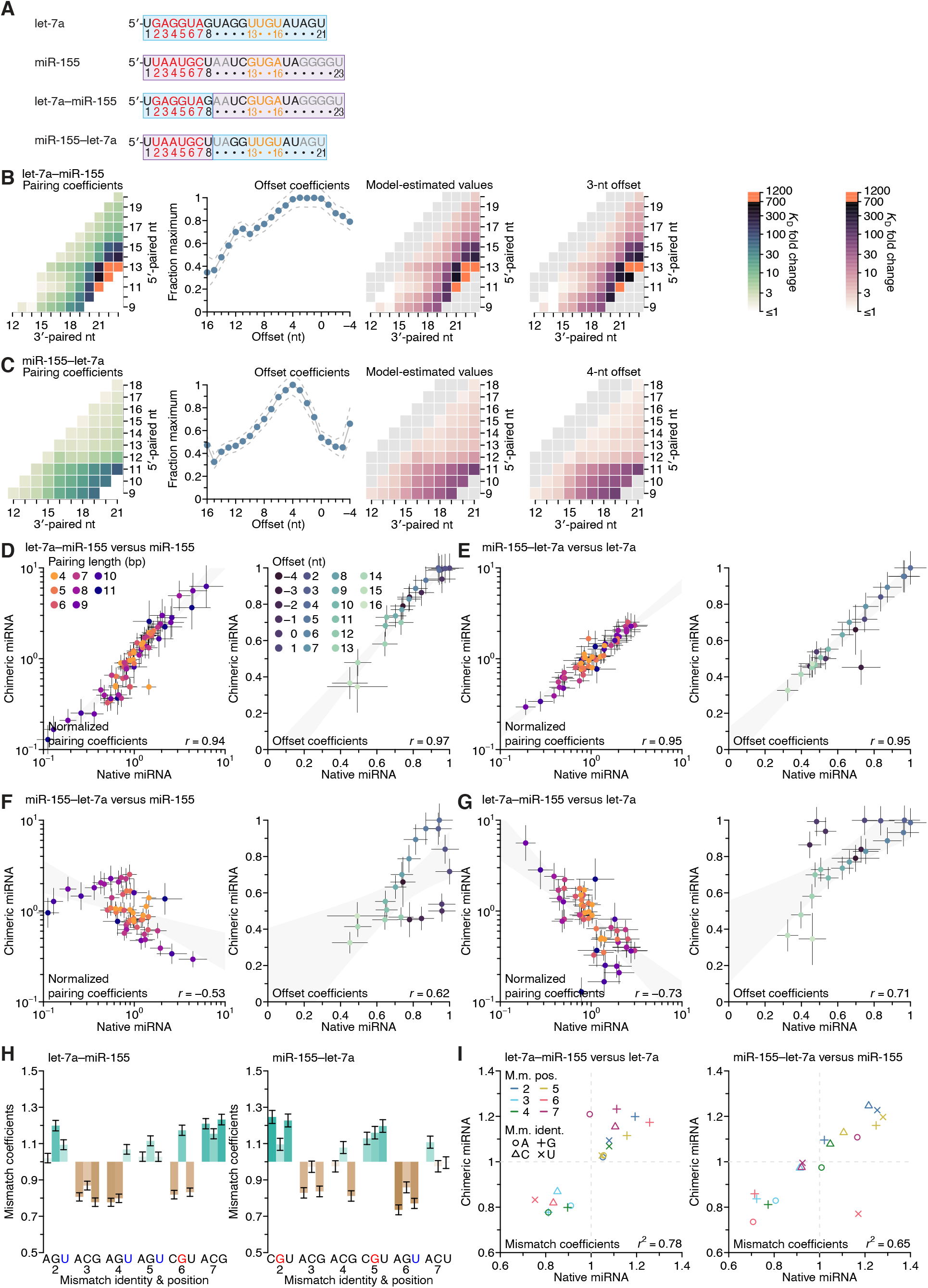
Independence of seed-mismatch and 3′-sequence effects. (**A**) Sequences of native let-7a, native miR-155, a chimeric miRNA containing the seed region of let-7a appended to nucleotides 9–23 of miR-155 (let-7a–miR-155), and a chimeric miRNA containing the seed region of miR-155 appended to nucleotides 9–21 of let-7a (miR-155–let-7a). (**B** and **C**) Pairing and offset coefficients describing the 3′-pairing preferences of let-7a–miR-155 (**B**) and miR-155–let-7a (C). Orange cells indicate pairing coefficients or *K*_D_ fold-change values between 700–1200. Otherwise, this panel is as in Figure 4A. (**D**) Comparison of the pairing and offset coefficients determined for let-7a–miR-155 with those of miR-155. Left, comparison of pairing coefficients, after first dividing by the geometric mean of all pairing coefficients of the same length for that miRNA, which normalized the trivial effect of pairing length. Points are colored according to pairing length, as in Figure 2A; error bars indicate 95% confidence intervals. Right, comparison of offset coefficients, colored from light blue to dark blue, progressing from offsets of − 4 to +16 nt; error bars indicate 95% confidence intervals. For each graph, the gray region indicates the 95% confidence interval for the linear least-squares fit to the data (*r*, Pearson correlation coefficient). (**E**) Comparison of the pairing and offset coefficients determined for miR-155–let-7a with those of let-7a. Otherwise, this panel is as in (D). (**F**) Comparison of the pairing and offset coefficients determined for miR-155–let-7a with those of miR-155. Otherwise, this panel is as in (D). (**G**) Comparison of the pairing and offset coefficients determined for let-7a–miR-155 with those of let-7a. Otherwise, this panel is as in (D). (**H**) Seed-mismatch coefficients of the let-7a–miR155 (left) and miR-155–let-7a (right) chimeric miRNAs. Otherwise, this panel is as in Figure 4F. (**I**) Correspondence between mismatch coefficients of chimeric miRNAs and those of their seed-native miRNAs. For let-7a–miR-155 (left) and miR-155–let-7a (right), the values from (H) are plotted against those of Figure 4F (*r*^2^, coefficient of determination). Colors and symbols indicate the position and nucleotide identity, respectively, of each seed mismatch coefficient (key).

Both of the chimeric miRNAs had 3′-pairing and offset preferences that were remarkably similar to those of the natural miRNAs containing the same 3′ sequences (Figures 4A, 4C, 5B, and 5C). Indeed, comparison for each chimeric miRNA to its 3′-native sequence revealed a high correspondence of pairing and offset coefficients (Figures 5D and 5E), and comparison of each chimeric miRNA to its seed-native miRNAs showed much lower correspondence of the pairing and offset coefficients. (Figures 5F and 5G). Furthermore, the fitted slopes for four 3′-native comparisons approached unity (range, 0.80–1.17), which showed that the effect sizes of these preferences were similar regardless of whether the coefficients were derived from chimeric or native miRNA datasets.

When analyzing the influence of the 18 seed mismatches on the affinity of 3′ pairing, miRNAs with the same seed sequence but different 3′ sequences had largely similar preferences (Figures 4F, 5H, and 5I). The two most prominent exceptions were the increased affinity in the context of a mismatched A at position 7 of the let-7–miR-155 chimeric miRNA (Figure 5I, left), and the decreased affinity in the context of a mismatched U at position 6 of the miR-155–let-7a chimeric miRNA (Figure 5I, right). Despite these outliers, the influence of the seed mismatch on the magnitude of 3′-pairing affinity depended primarily on the seed-mismatch type and position, with relatively little dependence on the sequence of the 3′ region.

### Sequence preferences for 3′ sites are maintained at adjacent positions

We next investigated the extent to which the positional dependencies of 3′ pairing affinity were dependent on the positions themselves rather than the sequence at these positions. To do so, we generated 3′-pairing profiles of let-7a variants that shifted the native let-7a 3′ sequence by a single nucleotide in either direction (let-7a(− 1) and let-7a(+1)) while maintaining the miRNA length (Figure 6A). Comparison of the pairing preferences of let-7a(− 1) and let-7a(+1) showed corresponding shifts of the pairing coefficients, such that the characteristic benefit of pairing to the G found at nucleotide 11 of the native miRNA was maintained in both variants. Thus, the most consequential nucleotide shifted to 10 when this G shifted to position 10 in let-7a(− 1), and likewise, it shifted to 12 for let-7a(+1) (Figures 6B–6D).

**Figure 6.**
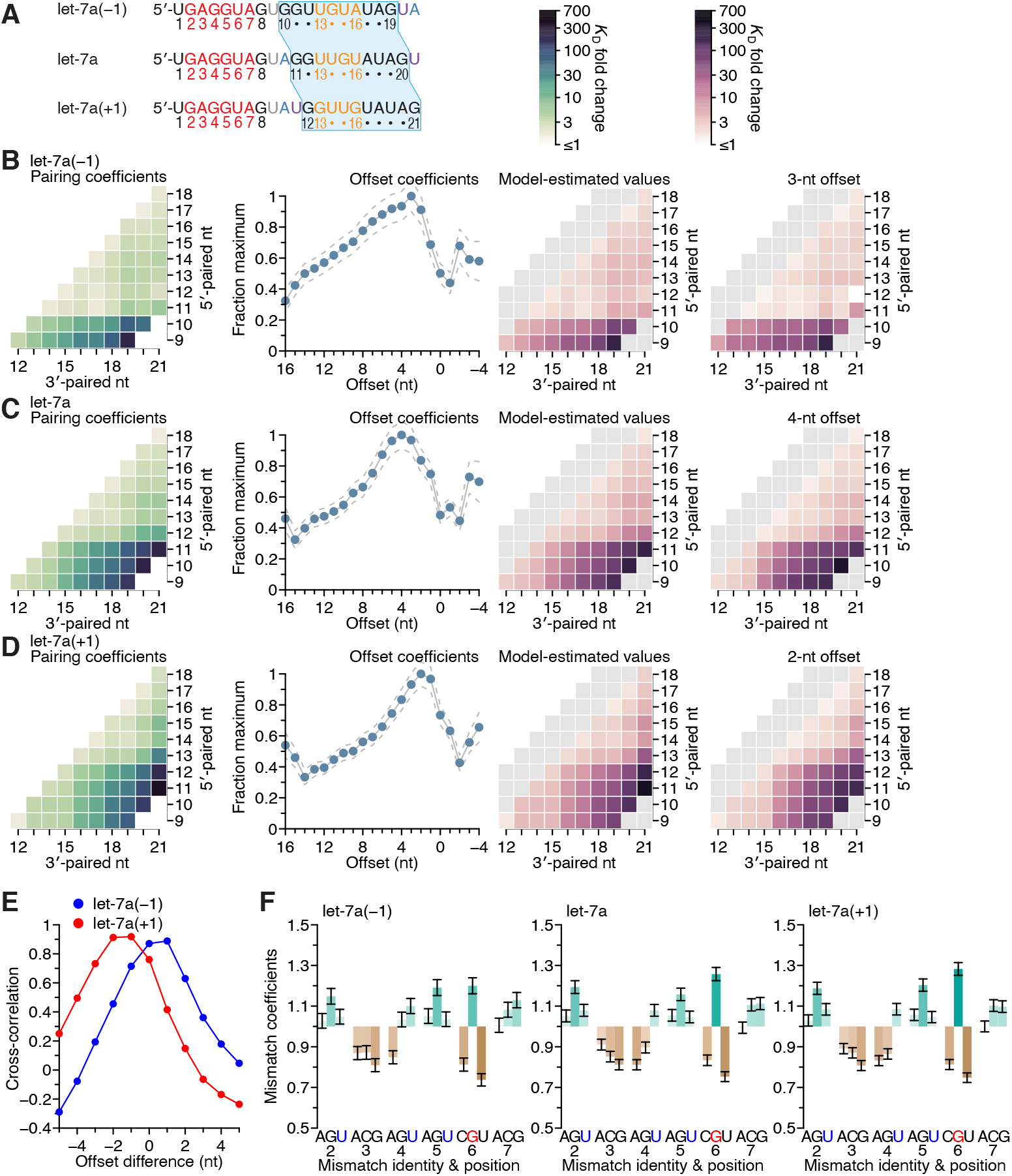
Sequence preferences for 3′ sites are maintained at adjacent positions. (**A**) Sequences of let-7a(− 1), which has a 3′ region permuted one nucleotide toward the 5′ end, native let-7a, and let-7a(+1), which has a 3′ region permuted one nucleotide toward the 3′ end. The 3′ sequence shared between all three miRNAs is shaded in blue, and the A and U nucleotides that were rearranged to generate the permuted variants are in blue and purple, respectively. (**B**–**D**) Pairing and offset coefficients describing the 3′-compensatory pairing of let-7a(− 1) (B), let-7a (C, redrawn from Figure 4A, for comparison), and let-7a(+1) (D). Otherwise, this panel is as in Figure 4A. (**E**) Cross-correlations of offset coefficients for either let-7a(− 1) (blue) or let-7a(+1) (red) with respect to those of let-7a (B and C, middle-left), plotted as a function of the difference in offset. (**F**) Effects of seed mismatches on 3′-pairing affinities of let-7a(− 1) (left), let-7a (middle, redrawn from Figure 4F, for comparison), and let-7a(+1) (right). Otherwise, this panel is as in Figure 4F.

The offset preference of let-7a(+1) was between +2 and +3 nt, which corresponded to a 1-to 2-nt shift compared to the +4-nt offset preference of let-7a (Figure 6E). These results supported the idea that fewer nucleotides were required to bridge the seed and 3′ pairing when the 3′ pairing started at position 12 rather than position 11. The offset preference of let-7a(− 1) shifted between 0 and +1 nt, which indicated that a target loop length of 6 nt was sufficient for inclusion of the centrally located GG dinucleotide within the 3′ pairing, regardless of whether this dinucleotide resided at positions 10 and 11 or at positions 11 and 12. Moreover, the seed mismatch preferences of both let-7a derivatives were nearly identical to those of the native let-7a (Figure 6F; *r*^2^ = 0.91 and 0.99 for let-7a(− 1) and let-7a(+1), respectively). Considered together, these results provided further evidence of the independent effects of the seed and 3′ region on 3′ pairing, with the behavior of the 3′ region depending on both position and sequence, with sequence preferences transferable to nearby positions, especially if compensating changes to the offset optimize the length of the target segment bridging the seed and 3′ pairing.

### Effects of mismatches within 3′ sites are consistent across miRNAs but explained poorly by the nearest-neighbor model

Having systematically analyzed the effects of seed-mismatch identity and of the length, position, and offset of perfect 3′ pairing, we next sought to measure the effects of any imperfections— i.e., mismatches, wobbles, or bulged nucleotides—within this 3′ pairing. Accordingly, we measured the affinities of variants of each site considered thus far, looking at each possible variant that had one of the eight possible imperfections at one position within the site. These eight imperfections considered at each position of interest included three possible mismatched nucleotides (including G:U wobbles), four possible single-nucleotide bulges (occurring opposite the linkage of two miRNA positions and assigned to the more 3′ miRNA position), and one single-nucleotide deletion (i.e., a bulged nucleotide in the miRNA). Consideration of these variants together with the original sites with perfect contiguous pairing resulted in the measurement of *K*_D_ values for 38,108 let-7a sites, 44,190 miR-1 sites, and 52,166 miR-155 sites. Analysis of these variants in the context of the best sites at each length (Figures 7A–7C and S12) revealed no imperfections that increased 3′-site affinity, which indicated that there were no positions at which the altered helical geometry of a mismatch was favored over Watson–Crick pairing. When comparing effects of internal mismatches to those of mismatches occurring at the end of the pairing, no striking differences were observed. Nonetheless, effects at some positions were more striking than others, with larger effects observed for mismatches involving any of nucleotides 11–15 of let-7a (Figure 7A), 12–15 of miR-1 (Figure 7B), and 15–22 of miR-155 (Figure 7C), which concurred with the importance of extending pairing to G11-G12, G12, and G19-G20-G21-G22 of the respective miRNAs.

**Figure 7.**
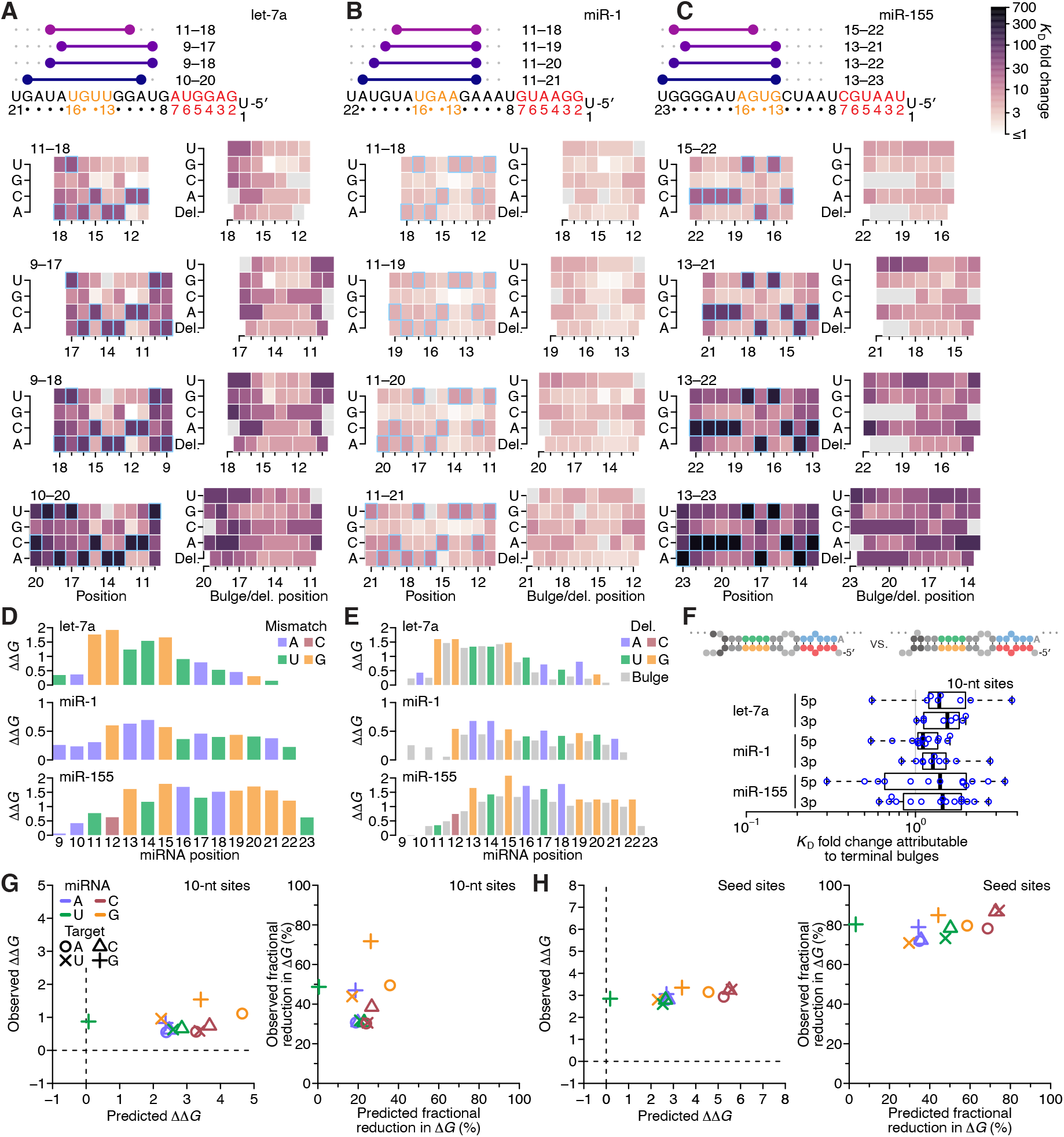
The impact of mismatched, bulged, and deleted target nucleotides on 3′-compensatory pairing. (**A**) The effect of mismatched, bulged, and deleted target nucleotides on 3′-compensatory pairing to let-7a. At the top is a schematic depicting the position of highest-affinity 3′-pairing for 3′ sites of lengths 8–11 nt, redrawn from Figure 2B. Below, at the left are heat maps corresponding to each of the pairing positions shown above, indicating the affinities with each of the four possible nucleotides at each position of the site. Cells corresponding to the Watson– Crick match are outlined in blue. Cells for affinities of mismatches that could not be calculated due to sequence similarity to another site type (e.g., the mismatched U across from position 14, which was indistinguishable from a 6mer-m8 seed site) are in gray. To the right are heat maps that correspond to the same pairing ranges but indicate the effects of a bulged or a deleted (del.) 3′-target nucleotide. A bulged nucleotide at position *n* corresponded to an extra target nucleotide inserted between the nucleotides pairing to miRNA positions *n* – 1 and *n*. (**B** and **C**) The effects of mismatched, bulged, and deleted target nucleotides on 3′-compensatory pairing to miR-1 (B) and miR-155 (C). Otherwise, these panels are as in (A). (**D**) Profiles of 3′-pairing mismatch tolerances. Each bar represents the ΔΔ*G* value when averaging over the three possible mismatches at that position, for let-7a (top), miR-1 (middle), and miR-155 (bottom). Each of the mismatch ΔΔ*G* values was an average of the values observed in the context of each 10-nt 3′ site that included the position. The color indicates whether the miRNA nucleotide was an A (blue), U (green), C (red), or G (yellow). (**E**) Profiles of tolerances to bulged and deleted nucleotides. Each colored bar represents the ΔΔ*G* value when deleting the target nucleotide complementary to the miRNA at that position, and each dark gray bar represents the ΔΔ*G* value when averaging all four bulged nucleotide possibilities occurring at that inter-nucleotide position, for let-7a (top), miR-1 (middle), and miR-155 (bottom). Each of the ΔΔ*G* values represents the average of the values observed in the context of each 10-nt 3′ site that included the position. The color of each bar corresponding to a deletion indicates whether the resulting bulged miRNA nucleotide was an A (blue), U (green), C (red), or G (yellow). (**F**) The tolerance of bulged nucleotides near the ends of 3′ sites. Plotted are ratios of *K*_D_ fold-changes comparing a site that has a bulged nucleotide between the penultimate and terminal base pairs with a site that does not have the terminal base pair (in which case, the bulged nucleotide in the former pairing architecture becomes a terminal mismatch). The box plots indicate the minimum, lower quartile, median, upper quartile, and maximum values. For each of the three miRNAs, comparisons are made for bulges occurring at the 5′ end of the 3′ pairing (5p), and at the 3′ end of the 3′ pairing (3p). The vertical gray line indicates a *K*_D_ fold-change ratio of 1.0. At the top is an example of a 3p comparison. (**G**) Comparison of the measured mismatch ΔΔ*G* values in 3′ sites with values predicted by nearest-neighbor rules. Left, comparison of the average measured ΔΔ*G* value with the average predicted value for each of the 12 possible miRNA–target mismatch combinations. Right, comparison of measured and predicted average fractional reduction in Δ*G* attributed to each mismatch. The fractional reduction was given by (Δ*G*_WC_ − Δ*G*_mm_)/Δ*G*_WC_, where Δ*G*_WC_ corresponds to the Δ*G* of the site with full Watson–Crick pairing, and Δ*G*_mm_ corresponds to the Δ*G* of a site containing the mismatch. These average values were calculated using *K*_D_ fold-change values determined for 10-nt sites, first averaging results for the same position over all 10-nt sites that included the position, then averaging results for that mismatch across all positions of the miRNA that had that mismatch, and then averaging the results across all three miRNAs. Colors and symbols indicate miRNA and target nucleotide identities, respectively (key). (**H**) Comparison of the measured seed mismatch ΔΔ*G* values with values predicted by nearest-neighbor rules. For each mismatch type, both the measured and predicted ΔΔ*G* values were the average over all occurrences within positions 2–7 for let-7a, miR-1, miR-155, miR-124, lsy-6, and miR-7, using relative *K*_D_ values from analyses of random-sequence AGO-RBNS results. Otherwise, this panel is as in (G).

To investigate mismatch tolerance across the range of miRNA 3′-end positions, we calculated the geometric mean of the *K*_D_ fold change for a mismatch at each position for all three miRNAs, averaging both over the three mismatches at each position and over each of the 10-nt sites that contained the position (Figure 7D). As expected, reduced binding affinity tracked with the importance of the positions for 3′ pairing, with greatest effects observed at G11 and G12 of let-7a, the G12–G15 of miR-1, and G13 and G15–G21 of miR-155 (Figure 7D). The greater importance of pairing to G13 compared to pairing to C12 of miR-155 further supported the idea that pairing to G had a greater impact over pairing to C in the miRNA 3′ region. Nonetheless, extending the analyses of mismatches, wobbles, and bulges to the random-sequence RBNS datasets previously acquired for six miRNAs (Figure S13) indicated that disrupting pairing to either C13 or C15 of the C13-G14-C15 trinucleotide of lsy-6 greatly reduced affinity. Thus, in some nearest-neighbor and positional contexts, pairing to a miRNA C nucleotide can be as important as pairing to a miRNA G nucleotide. More generally, these results showed that the effect of a mismatch to a particular nucleotide was informed primarily by the overall importance of that miRNA nucleotide for pairing (as determined by its nucleotide identity and position within the miRNA 3′ end), irrespective of whether the target nucleotide fell within the middle or terminus of the 3′ site.

To summarize the positional tolerance of bulges, we averaged the effects of the four bulges at each inter-nucleotide position and over each of the 10-nt sites containing that position. Likewise for the deletions, we averaged each single possibility over the 10-nt sites containing that position (Figure 7E). At each position, the severity of both types of lesions tracked with that observed for the mismatches, with the effects of deletions generally similar to those of their corresponding mismatches, and effects of bulged nucleotides marginally less severe.

To examine if the benefit of bulged nucleotides over mismatched nucleotides applied to the very 5′ and 3′ ends of 3′ sites, we considered all possible 10-nt sites for all three miRNAs with programmed libraries, and calculated the fold difference in relative *K*_D_ observed when comparing a site with a terminal mismatch to that of the site with a corresponding terminal bulged nucleotide (i.e., the site variant in which the target nucleotide following the mismatch can pair to the mismatched miRNA nucleotide). For each miRNA, a small but significant benefit to terminal bulges was observed (Figure 7F; *p* = 2.4 × 10^− 5^, 1.4 × 10^− 6^, and 4.5 × 10^− 4^ for let-7a, miR-1, and miR-155, respectively; one-tailed Wilcoxon signed rank test). Thus, an isolated complementary target nucleotide separated from a longer contiguous stretch of pairing can contribute modestly to site affinity.

To enable comparison of the observed effects of mismatches with those predicted by the nearest-neighbor model of RNA duplex stability, we calculated the ΔΔ*G* of each mismatch in the context of all 10-nt 3′ sites of the three miRNAs. We first averaged these values over all the contiguous sites, and then over all positions with the same miRNA nucleotide, and then over the three miRNAs, resulting in one global average ΔΔ*G* value for each of the 12 possible miRNA– target mismatch possibilities. Comparison of these values with those predicted using the nearest-neighbor parameters revealed that the effects of the mismatches were typically much lower than expected for RNA in solution, with no strong relationship between the observed and predicted ΔΔ*G* values (Figure 7G, left; *r*^2^ = 0.02). The outlier in this analysis was the miRNA– target U:G wobble, which was as disruptive as the typical mismatch but predicted to be much less so (Figure 7G, left, green +). Next, to account directly for the reduced binding energy of the fully complementary sites in comparison to their predicted Δ*G* values, we compared the average observed and predicted fractional reduction in Δ*G* of each site caused by each of the twelve mismatch values (Figure 7G, right). For eight of 12 mismatches, the fractional reduction in Δ*G* was within 10% of its prediction, but the miRNA–target A:G, G:G, G:U, and U:G mismatches respectively caused 31%, 42%, 21%, and 48% more reduction in binding energy than predicted. These results indicated that the nearest-neighbor parameters were not suited for predicting the contribution of miRNA 3′ pairing in three respects: 1) the overall contribution to binding energy was far less than that predicted, 2) mismatched target G nucleotides were relatively more deleterious than predicted, and 3) wobble pairing was relatively less favorable than predicted. Indeed, the U:G possibility, which both contained a target G nucleotide and was a wobble, was the mismatch with the greatest deviation from expectation.

For comparison, we repeated these analyses for mismatches to the miRNA seed (i.e., miRNA positions 2–7) within the context of canonical 8mer pairing, calculating the average ΔΔ*G* and the fractional reduction in Δ*G* for each type of mismatch for each of the six miRNAs for which there was random-sequence RBNS data (McGeary et al., 2019) (Figure 7H). These analyses indicated that the effects of mismatches within seed pairing also did not agree with predicted pairing energetics, albeit differently than the effects of mismatches within 3′ pairing. First, a mismatch within the seed pairing had a much larger influence on ΔΔ*G* than did a mismatch within the 3′ pairing. Moreover, the reductions in binding affinities for mismatches within the seed pairing were even more regular than those for mismatches within the 3′ pairing, with a ∼3 kcal/mol detriment for each of the 12 mismatch/wobble possibilities (Figure 7H, left). The fractional reduction in Δ*G* had a similarly large and uniform effect size, with no subset of the mismatch possibilities showing a relationship with that predicted (Figure 7H, right). Thus, the binding preferences at both the seed and 3′ regions of the miRNA were not well characterized by nearest-neighbor rules, although the nature of the deviations differed in these two regions.

## DISCUSSION

An AGO-loaded miRNA can be divided into three regions: the seed region (nucleotides 2–8), the central region (nucleotides 9–10 or 9–11), and the 3′ region (Figure 1A) (Bartel, 2018). Because the most effective 3′ pairing is reported to center on nucleotides 13–16 (Grimson et al., 2007), some subdivide the 3′ region into the 3′-supplementary region (nucleotides 13–16), and the tail (nucleotides 17 to the terminus), while expanding the central region to include nucleotide 12 (Salomon et al., 2015; Schirle et al., 2014; Sheu-Gruttadauria et al., 2019b; Wee et al., 2012). The structure of AGO2–miR-122 bound to a 3′-supplementary site, which shows that miRNA nucleotides 9–11 are not available for pairing due to both helical distortion and inaccessibility caused by residues of the PIWI and L2 loop, seems to support the notion of a 3′-supplementary region at nucleotides 13–16 (Sheu-Gruttadauria et al., 2019b). However, greater affinities are observed with more extended 3′ pairing (Becker et al., 2019; Sheu-Gruttadauria et al., 2019a), and we found that 3′-site affinities nearly always increased as potential for pairing expanded to include most of the 3′ region—and in the positive-offset binding mode, some of the central region. Thus, productive 3′ pairing can encompass the entire miRNA 3′ region and should not be thought of as limited to a short 3′-supplementary region. Indeed, the study reporting that pairing to nucleotides 13–16 is most effective for supplementing seed pairing uses a model for predicting the efficacy of 3′ pairing that rewards extension of that pairing into the remainder of the 3′ region (Grimson et al., 2007).

Also problematic for the notion of a short 3′-supplementary region common to all miRNAs was our observation that the positions most important for 3′ pairing differed between different miRNAs. For example, at their optimal offsets, both let-7a and miR-124 preferred pairing to nucleotides 11–14 over pairing to nucleotides 13–16 (Figures 2B, 4A, S6A, and S6D), and the synthetic let-7a(− 1) preferred pairing to nucleotides 10–13 over pairing to nucleotides 13–16 (Figure 6B). Moreover, although miR-155 preferred pairing to nucleotides 13–16 over other 4-nt possibilities, when examining 7-nt 3′ sites, it preferred pairing to nucleotides 15–21 over pairing to nucleotides 13–16 (Figures 3C and 4B). These observations showing that the preferred positions of 3′ pairing can vary so widely between miRNAs, to include virtually any nucleotide downstream of the seed, argued strongly against assigning the same short 3′-supplementary region to all miRNAs.

Our observation that let-7a preferred pairing to nucleotides 11–14 over pairing to nucleotides 13–16 concurs with recent analyses of the relative importance of these nucleotides in *Caenorhabditis elegans. C. elegans* requires *let-7* repression of *lin-41* for viability, and this repression occurs through two 3′-compensatory sites that each have pairing to nucleotides 11– 19 of the miRNA (Aeschimann et al., 2019; Pasquinelli et al., 2000; Reinhart et al., 2000). Mutagenesis of individual nucleotides of the *let-7* miRNA indicates that nucleotides 11, 12, and 13 are each critical for viability, whereas nucleotides 14, 15, and 16 each have intermediate importance, and nucleotides 17, 18, and 19 each have no detectable importance (Duan et al., 2021). Inspection of our data for let-7a, examining the effects of mismatches within the 3′ site that has the same architecture as that of the two sites within *lin-41* (i.e., 9 bp of pairing beginning at position 11 with an offset of +1 nt) revealed a similar polarity, with mismatches near position 11 tending to be most consequential and those near position 19 tending to be least consequential (Figure S12D).

Although our results showed that preferred pairing often did not correspond precisely to positions 13–16, preferred pairing did always at least partially overlap this segment. Moreover, as pairing lengths increased from 4 to 6 bp, overlap between preferred pairing and this segment increased, such that the preferred 6-nt sites for let-7a, miR-1, miR-155, miR-124, miR-7 and lsy-6 each included pairing to miRNA nucleotides 13–16. The only exception we observed was the preferred 6-nt site for synthetic let-7a(− 1), which paired to nucleotides 10–15. Thus, our results explain why an overall preference for pairing to nucleotides 13–16 was detected in meta-analyses of both functional data for 11 miRNAs and evolutionary conservation of sites for 73 miRNA families (Grimson et al., 2007). Our key added insight is that sequence identity in the 3′ region—particularly the placement of stretches of G residues—imparts additional preferences that supplement the positional preferences to specify different optimal regions of 3′ pairing for different miRNAs.

Another key insight is evidence of two distinct 3′-binding modes, observed as different offset preferences of let-7a, miR-124, lsy-6, and miR-7 with and without pairing to nucleotide 11 (Figures 2B, 2C, and S7). In one binding mode, an offset of 0 nt is optimal for 3′ pairing starting at position 12, whereas in the other binding mode, additional nucleotides are required to bridge pairing to positions 10 or 11, resulting in optimal offsets that exceed 0 nt. In a crystal structure of AGO2–miR-122 bound to a 3′-supplementary target that pairs to nucleotides 13–16 with an offset of 0 nt, nucleotide 12 is the first nucleotide available for pairing, whereas pairing to nucleotide 11 is occluded by the central gate (Sheu-Gruttadauria et al., 2019b). We suggest that this structure reflects the conformation of the zero-offset binding mode, as it provides a physical model for why extension of potential pairing from nucleotide 12 to 11 results in almost no increased binding affinity for sites with an offset of 0 nt (Figures S7B, S7D, S7F, and S7G).

However, another structure will be required to visualize the positive-offset binding mode that enables optimal pairing to let-7a and miR-124, as well as strong pairing to lsy-6 and miR-7. Genetically identified sites inferred to be utilizing this second binding mode include the two let-7a sites within the 3′ UTR of *C. elegans lin-41*, which both include pairing to nucleotide 11 and an offset of +1 nt, as well as the first lsy-6 site within the 3′ UTR of *C. elegans cog-1*, which includes pairing to nucleotide 11 and an offset of +2 nt. The discovery of these two binding modes required knowledge of the interplay between preferred pairing position and preferred pairing offset, which underscored the utility of obtaining affinity measurements for a large diversity of 3′ sites.

The length of a miRNA can modulate its 3′-pairing affinity, in that a 23-nt derivative of miR-122 has a 3-fold longer dwell time than its 22-nt counterpart (Sheu-Gruttadauria et al., 2019b). Of the miRNAs that we examined, miR-155 and miR-7 were each 23 nt in length, whereas the others were shorter. These two miRNAs had the strongest and the weakest 3′ pairing, respectively. The weak 3′ pairing of miR-7 indicated that although increased miRNA length can sometimes improve 3′ binding affinity, it cannot substitute for other features required for high affinity to the miRNA 3′ region.

Early attempts to either explain targeting efficacy or predict target sites used scores incorporating, among other things, the predicted binding energy between the miRNAs and their proposed targets (Doench and Sharp, 2004; Enright et al., 2003; Krek et al., 2005; Lewis et al., 2003; Rajewsky and Socci, 2004). That these metrics were less useful in identifying consequential 3′ pairing than simpler rubrics scoring only the length and position of complementarity (Grimson et al., 2007) suggests that the parameters derived from interactions of purified RNAs in solution are not directly relevant to miRNAs associated with AGO. The breadth of our affinity measurements provided the ability to assess why such parameters are not as useful. Although high correspondence was observed between the predicted Δ*G* and measured 3′ pairing affinities (Figure S5A), for miR-1 this relationship nearly disappeared when normalizing for pairing length (Figure 4E). For let-7a and miR-155 a relationship was retained after normalizing for length, but four factors limit the utility of using this relationship for ranking target predictions. The first is the strong effect of position, with complementarity to the seed much more consequential than complementarity to the 3′ region, and complementarity at some positions in the 3′ region more consequential than complementarity to others, and much more consequential than complementarity to positions 1, 9, and often, 10. The second is the effect of primary sequence, as illustrated by the outsized benefit pairing to the G11, G12, and G20 nucleotides of let-7a, miR-1, and miR-155, respectively (Figures S5B and S5C). The third is the poor relationship between the predicted and measured effects of some internal mismatches and wobbles (Figure 7G), and the fourth is a lack of a consistent relationship between predicted Δ*G* and measured binding affinities between miRNAs (Figure S5A, comparing the slope for miR-1 with that of either let-7a or miR-155).

Comparison of the 3′ regions of the four miRNAs that were more effective at 3′ pairing with those of the two that were not suggested a feature that might have conferred higher 3′-pairing affinity: the presence of two or more adjacent G nucleotides (e.g., the G11-G12 of both let-7a and miR-124, and the G19-G20-G21-G22 of miR-155). Although lsy-6 did not have an oligo(G) stretch, it did have a well-positioned C13-G14-C15 trinucleotide, which together with G11 was critical for pairing affinity. When considering all four miRNAs together, as well as the lack of any GG, CG, or GC dinucleotides within the 3′ regions of miR-1 or miR-7, we suggest that miRNAs with GG, CG, or GC dinucleotides within positions 13–16 are the ones most likely to participate in productive 3′ pairing, and that pairing that extends to an oligo(G) sequence outside of positions 13–16 will preferentially enhance affinity.

The importance of pairing to miRNA G nucleotides, and not C nucleotides (other than the C13-G14-C15 of lsy-6), suggested that a miRNA–target G:C base pair is read out differently than a C:G base pair. Perhaps G nucleotides participate in base-stacking interactions that position or pre-organize the guide strand to favor nucleation of 3′ pairing. Alternatively, the explanation might involve target-site accessibility. Pairing to a C in the miRNA 3′ region would require a G in the vicinity of the seed match, which compared to a C would cause poorer target-site accessibility (McGeary et al., 2019), thereby reducing the net contribution to binding.

Our results also revealed a functional difference between 3′-supplementary and 3′-compensatory pairing. The added affinity of a 3′ site was relatively constant when it supplemented different sites that had seed matches (Figures 4H and S11), whereas it varied in the context of different 3′-compensatory sites that had different seed mismatches (Figures 4F and S8–S10). The effects of seed mismatches were miRNA-specific and unrelated to their binding affinities (Figure 4G). Additionally, our experiments using chimeric miRNAs demonstrated the separability of the mismatch effects from the length, position, offset, and nucleotide-identity preferences of the 3′ region (Figure 5).

Pairing to the miRNA 3′ region not only increases site affinity and target repression, but it can also influence the stability of the miRNA itself, in a process called target-directed miRNA degradation (TDMD) (Ameres et al., 2010; Bitetti et al., 2018; Cazalla et al., 2010; Kleaveland et al., 2018; Mata et al., 2015). The handful of target sites known to trigger TDMD have diverse 3′-pairing architectures. For example, degradation of miR-7 triggered by the cellular Cyrano transcript occurs through a canonical 8mer site supplemented with a 3′ site with 14 contiguous pairs to the 3′ end of the miRNA (Kleaveland et al., 2018), whereas degradation of miR-27a triggered by the m169 RNA from murine cytomegalovirus occurs through a canonical 7mer-A1 site supplemented with a 3′ site with only six contiguous pairs to the 3′ end of the miRNA (Marcinowski et al., 2012). Our finding that miR-7 has the weakest 3′ pairing among the six miRNAs we studied provides a potential explanation as to why its TDMD trigger Cyrano has such a long 3′ site.

The crystal structures of several known TDMD substrates bound to their corresponding TDMD-inducing target sites reveal a distinct conformation for these AGO–miRNA–target RNA ternary complexes in comparison to ternary complexes that have supplementary pairing involving only nucleotides 13–16 (Sheu-Gruttadauria et al., 2019b, 2019a). During TDMD, this distinct conformation is thought to be recognized by the ZSWIM8 E3 ubiquitin ligase, causing AGO proteolysis through the ubiquitin–proteasome system, which exposes the miRNA to degradation by cellular nucleases (Han et al., 2020; Shi et al., 2020). Our discovery of the two 3′ binding modes raises the question of whether one of them might be more compatible with TDMD, perhaps due to a preference of the ZSWIM8 E3 ligase. Although the TDMD ternary complexes of the published structures all have 3′ pairing beginning at nucleotide 12 or later and offsets of 0 or − 1 nt (Sheu-Gruttadauria et al., 2019a) and thereby represent the zero-offset binding mode, the 3′ pairing between miR-7 and Cyrano begins at G11 and has a +2-nt offset, which represents the positive-offset binding mode. Thus, the two 3′ binding modes both appear to be compatible with either of the two gene-regulatory processes that involve 3′ pairing— TDMD and miRNA-mediated repression.

## ACKNOWLEDGEMENTS

We thank K. Lin and T. Pham for helpful discussions, the Whitehead Genome Technology Core for high-throughput sequencing, and members of the Bartel lab for comments on this manuscript. This work was supported by NIH grants GM118135 (D.P.B.) and GM123719 (N.B.).

D.P.B. is an investigator of the Howard Hughes Medical Institute.

## AUTHOR CONTRIBUTIONS

N.B. performed the experiments and the initial analyses, with help from S.E.M. S.E.M. performed the final analyses. S.E.M., N.B., and D.P.B. designed the study and wrote the manuscript.

