## Supplementary material for "Pairing to the microRNA 3′ region occurs through two alternative binding modes, with affinity shaped by nucleotide identity as well as pairing position"

**This PDF file includes:**

Experimental Materials and Methods
Computational and Mathematical Methods
Supplementary Figures S1–S13

**Other Supplementary Materials for this manuscript include the following:**

Table S1: Oligonucleotides used in this study (Excel file)

### EXPERIMENTAL MATERIALS AND METHODS

#### Human Embryonic Kidney 293T (HEK293T) Cells

HEK293T cells were cultured in DMEM (VWR) with 10% fetal bovine serum (Clonetech) at 37°C with 5% $CO_2$, and split every third day at ~90% confluency.

#### Purification of AGO2–miRNA complexes

AGO2–miRNA complexes were generated and purified as described (McGeary et al., 2019).

#### Preparation of programmed RNA libraries

For each miRNA, a programmed library was constructed by in vitro transcription of multiple chemically synthesized DNA libraries, and then mixing these RNA libraries after gel purification. Each library contained 25 nt of entirely randomized sequence, followed by an 8-nt programmed site, followed by either 5 nt of random sequence (let-7a and miR-1 libraries) or 4 nt of random sequence (miR-155 library). For every AGO-RBNS experiment other than that with native miR-155, the final library was made by mixing six different libraries, in which each of the six libraries had at its programmed site an 8mer containing a mismatch to one of the six seed nucleotides. For the experiment with native miR-155, the library was assembled by mixing the six 8mer-mismatch libraries with a seventh library that had a 6mer at the programed site, flanked by non-A residues at positions 1 and 8.

Each individual library was transcribed from synthetic DNA (IDT) and purified as described (McGeary et al., 2019). The fraction of reads representing each of the 18 seed-mismatch sites, expected to be 5.6%, was measured by sequencing of each input library and found to be 3.4–8.0% for let-7a rep. 1, 2.9–8.7% for let-7a rep. 2, 3.3–7.8% for miR-1, 2.2–6.3% for miR-155, 3.4–8.0% for let-7a(+1) and let-7a(−1), 3.3–7.6% for let-7a–miR-155, and 2.6–6.1% for miR-155–let-7a.

#### AGO-RBNS

AGO-RBNS was performed as described (McGeary et al., 2019).

### COMPUTATIONAL AND MATHEMATICAL METHODS

#### Analysis of *k*-mer enrichments

Positional enrichments of 8-nt *k*-mers were calculated by comparing the reads from RNA bound to 840 pM AGO2–let-7a complex to reads from the input library. For each of the two sets of reads, reads that contained one of the 18 possible 8mer-mismatch sites in the correct position (such that the CUACCUCA 8mer-consensus sequence spanned positions 26–33 of the read), but did not contain a canonical 8mer, 7mer-m8, 7mer-A1, 6mer, 6mer-m8, or 6mer-A1 site, were used to enumerate all possible 8-nt *k*-mers at each position within the library. Both count tables were normalized such that they summed to 1, and the normalized count table corresponding to the bound sample was divided by that of the input library to arrive at the enrichment of each *k*-mer at each position within the library. These *k*-mers were ranked according to the sum of the top five positional enrichments of each, considering each possible 8-nt window located entirely within the random sequence region 5′ of the programmed site. These 8-nt windows are labeled in Figure 1D relative to the position in the programmed site designed to pair to miRNA position 8, such that *k*-mer position 9 of Figure 1D corresponds to positions 18–25 when counting from the 5′ end of the random sequence region, and *k*-mer position 26 corresponds to library positions 1–8.

**Assignment of miRNA sites within reads from the programmed libraries**

When counting seed sites and contiguously paired 3′ sites within the random-sequence region of the programmed libraries, reads were first queried for whether they included a canonical or 8mer-mismatch site at the programmed region (i.e., at library positions 26–33, counting from the 5′ end of the random-sequence region). Reads containing a canonical site (e.g., an 8mer or 7mer-m8 site) in the programmed region despite its design were still counted and used for relative $K_D$ assignment, but their measured relative $K_D$ values were not used in any further analyses, owing to the ambiguity of whether the presence of these reads was the result of errors during chemical synthesis, in vitro RNA transcription, library preparation, or sequencing.

Those reads containing a seed site (defined as an 8mer, 7mer-m8, 7mer-A1, 6mer, 6mer-m8, or 6mer-A1 site, or one of the 18 possible canonical 8mer sites with a mismatch within positions 2–7) at the programmed region were further assigned to one of four categories based on whether the random region contained: 1) neither a seed site nor a 3′ site (defined as a site of 4–11 nt of contiguous complementarity to a region of the miRNA spanning position 9 to the 3′-most nucleotide), 2) at least one seed site but no 3′ sites, 3) at least one 3′ site but no seed sites, or 4) at least one seed site and at least one 3′ site. This categorization enabled read enrichment from category 3 to be compared with that of category 1 as an indication of the contribution of pairing to the 3′ end with minimal influence from seed binding outside the programmed region.

The categorization of each read proceeded as follows: 1) any canonical or 8mer-mismatch sites other than the programmed site were identified, looking within the entire random-programmed-random region with three nucleotides of constant sequence appended to the 5′ and 3′ ends of the read, 2) the 28-nt segment spanning library positions 9–37 (which included three nucleotides of the 5′ constant sequence) was queried for the longest contiguous match to the miRNA 3′ end, retaining multiple sites in the case of ties. Any site(s) 4–11 nt in length that was not contained within a canonical or 8mer-mismach site found within this 28-nt segment was counted as a 3′ site, and 3) the read was then assigned to one of the four categories described above. In the case of a read with either a single 3′ site or a single seed site within the random region, the read was assigned to that site, recording the type of site, the identity of the programmed site, and the distance between the two. In the case of a read with either multiple 3′ sites or multiple seed sites within the random region, the read count was divided equally among the sites. In the case of a read with both a 3′ site(s) and a seed site(s), the read was split between all seed-and-3′-site pairs, recording the names of the seed, 3′, and programmed site for each.

This categorization yielded tables of counts for reads with 1) only programmed sites, 2) seed-and-programmed site pairs with positional information, 3) 3′-and-programmed site pairs with positional information, 4) seed-and-3′-and-programmed site triples with no positional information, and 5) reads with neither a canonical site nor an 8mer-mismatch site at the programmed site position. These tables were either used directly for relative $K_D$ estimation, or first pooled such that the identity of their programmed site was not considered. When pooling, all counts corresponding to reads with identical site-and-positional information were summed into two categories: those for which the programmed site was an 8mer-mismatch site, and those for which the programmed site was one of the canonical sites.

**Assignment of miRNA sites within reads from the random-sequence libraries**

When counting seed sites (including 8mer-mismatch sites) and contiguously paired 3′ sites within the random-sequence libraries, the 37-nt random-sequence region of each read was appended with 3 nt of constant sequence at each end, except in the case of miR-1, for which the 5′-most 36 nt of the random-sequence region was appended with 3 nt of only the 5′ constant sequence, due to the sequence bias present at the very 3′ end of these libraries caused by erroneous lack of a TCG sequence in the 3′ constant region required for pairing to the Illumina reverse-primer sequence during bridge-amplification (McGeary et al., 2019). The relevant portion of each read was queried for all seed sites (including 8mer-mismatch sites) and all 3′ sites between 4–11 nt in length, allowing individual seed sites to overlap, and individual 3′ sites to overlap. If the read contained (other than the programmed site) only seed sites or only 3′ sites, the read counts were split evenly between each site. If a read contained at least one seed site and at least one 3′ site, and if the 3′ site(s) fell upstream of a seed site, the 3′ site(s) was trimmed to remove any overlap with the seed site(s), and retained if the 3′ site(s) was ≥4 nt in length.

If one or more 3′ site persisted for the read after accounting for any seed-site overlap, the 3′ site(s) of length equal to the longest 3′ site(s) was retained. All possible bipartite sites associated with that read were then enumerated, in which each seed site was considered to form a bipartite site with all 3′ sites that were 5′ of that seed site. The read was then split among any bipartite sites identified. In the event that no bipartite sites were identified, the read was split among all the seed and 3′ sites equally. While this procedure ensured equal partitioning of read counts in cases of multiple seed and 3′ sites within a given read, in practice only a small fraction of reads contained multiple seed sites and a 3′ site, or a seed site and multiple 3′ sites.

This categorization yielded tables of counts for reads with 1) only seed sites, 2) only 3′ sites 3) seed-and-3′ bipartite sites with recorded inter-site spacing, and 4) neither a seed nor a 3′ site (referred to as “no site”). These tables were either used directly for relative $K_D$ estimation, or the bipartite sites were pooled to combine data for 3′ sites with different canonical sites or 8mer-mismatch sites prior to relative $K_D$ estimation. When pooling, all counts corresponding to reads with the same 3′ site and distance from the miRNA position 8 of the seed site were summed into two categories: those for which the seed site was an 8mer-mismatch site, and those for which the seed site was one of the canonical sites.

**Calculation of relative $K_D$ values**

Relative $K_D$ values were calculated as described (McGeary et al., 2019).

**Correction of relative $K_D$ values using data from random-library experiments**

Due to the deviation from the expected linear relationship between the relative $K_D$ values calculated for canonical sites (8mer, 7mer-m8, 7mer-A1, 6mer, 6mer-m8, and 6mer-A1 sites) and 8mer-mismatch sites (Figures S1B, left) we applied locally estimated scatterplot smoothing (LOESS) to generate an empirical correction to apply to these data, as has been done in analyses of metabolic-labeling data to correct mRNA abundance as a function of uridine content

(Schwanhäusser et al., 2011). The relative $K_D$ values of the canonical sites for the programmed-library experiments used for this correction were derived from geometric mean of the relative $K_D$ values of each site measured at each position at which they occurred within the library and in the context of each of the 18 programmed 8mer-mismatch sites, whereas the relative $K_D$ values of the 8mer-mismatch sites were derived from the reads associated with their occurrence at the programmed site in the absence of any seed sites or 3′ sites 4–11 nt in length in the random region. The relative $K_D$ values of these sites for the random-library experiments were derived from counts corresponding to instances of these sites within the reads.

We corrected the programmed-library relative $K_D$ values by calculating $R_i$, defined as:

$$R_i = \ln \frac{K_{r,i}}{K_{p,i}} \,, \tag{1}$$

where $K_{r,i}$ and $K_{p,i}$ refer to the relative $K_D$ values derived from the random-library and programmed-library experiments, respectively, for each site *i*. LOESS was used to fit a nonlinear function describing $R_i$ as a function of $K_{p,i}$:

$$R(K_p) \sim f_{LOESS}(x = \ln K_p) \,. \tag{2}$$

This function was then used to correct each programmed library–derived relative $K_D$ value by multiplying each value by the output of the function with itself as input:

$$K'_{p,i} \equiv K_{p,i} \times e^{f_{LOESS}(x = \ln K_{p,i})} \,. \tag{3}$$

The transformed $K'_{p,i}$ values were used throughout the study other than in Figure S1A and the left-hand panels of Figure S1B. LOESS was implemented in R using the *loess* function as part of the *stats* package, with “span” and “surface” parameters set to 10 and “direct”, respectively.

**Analysis of results from Becker et al. (2019)**

The 22,300 $K_D$ values measured for let-7a were inspected using the table provided as supplementary data (Becker et al., 2019). For each $K_D$ value, the target sequence was queried for a canonical or 8mer-mismatch site, hierarchically looking first for the 8mer, any 8mer-mismatch sites, and then any 7mer-m8, 7mer-A1, 6mer, 6mer-m8, or 6mer-A1 sites. If one or more of these sites were found, the target sequence 5′ of each site was queried for its longest stretch of complementarity to the 3′ region of let-7a (i.e., all nucleotides 3′ of position 8), and if this complementarity was between 4–11 nt in length, the $K_D$ value of that target sequence was assigned to that bipartite site, specifying its seed site, 3′ site, and intervening nucleotide length. If multiple 3′ sites were tied for the longest length, both 3′ sites were ascribed to the target sequence. Upon using the sequences to define the bipartite site information for each target RNA, only those target RNAs with a single bipartite site, or a single seed site and no 3′ pairing between 4–11 nt in length, were included in the downstream analyses. From these sites, we calculated the $K_D$ fold change for each available 3′-site, offset, and seed-site combination (Figure S3) by dividing the geometric mean of the $K_D$ values of target RNAs containing that bipartite site by the geometric mean of that of the target RNAs containing the seed site with no 3′ pairing, except when calculating the $K_D$ fold-change values for bipartite sites with the 8mer-xA5 seed site (Figure S3F). Because no target RNAs fit our criteria as containing only the 8mer-xA5 site and no 3′ pairing 4–11 nt in length, we used 10 nM as the reference $K_D$, which was the lower limit of detection measured, and was the measured $K_D$ for seven of the 16 8mer-mismatch sites present in the data.

**Calculation of pairing and offset coefficients**

In order to separate the intrinsic pairing preferences of each miRNA 3′ end from the effects of varying the offset of pairing, we fit a thermodynamic model of 3′-end binding efficacy to the $K_D$ fold-change values measured when summing the counts from each of the 18 8mer-mismatch sites. The model was constructed to produce a $\log_{10}$-tranformed $K_D$ fold-change value, denoted here using $\kappa$, as a function of the 5′ terminus of the pairing *i*, the 3′ terminus of pairing *j*, and the offset between the seed and 3′ pairing *k*. In order to make no assumptions regarding the thermodynamic nature of 3′-end binding (e.g., that each nucleotide would contribute independently to the binding energy), and as well to make no assumptions about the nature of the offset preferences, the model included two sets of categorical coefficients, one set $\alpha_{i,j}$ describing the 3′-pairing range as a function of the 5′ and 3′ termini of pairing indices *i* and *j*, and another set $\beta_k$ describing the offset preferences as a function of the offset index *k*:

$$\kappa(i,j,k) = \kappa\big(\alpha_{i,j}, \beta_k\big)\,. \tag{4}$$

Because the nature of the relationship between the pairing range and the offset preferences could not be known *a priori*, we constructed three variants of the model function $\kappa(i,j,k)$:

$$\kappa_a(i,j,k) = \alpha_{i,j} + \beta_k \tag{5.1}$$

$$\kappa_m(i,j,k) = \alpha_{i,j}\beta_k \tag{5.2}$$

$$\kappa_{mc}(i,j,k) = \alpha_{i,j}\beta_k + \gamma, \tag{5.3}$$

where $\kappa_a(i,j,k)$, $\kappa_m(i,j,k)$, and $\kappa_{mc}(i,j,k)$ describe additive, multiplicative, and multiplicative-plus-constant models. We note that an additive-plus-constant variant is trivially equivalent to an additive model, since the constant term can be subsumed by either of the $\alpha_{i,j}$ or $\beta_k$ coefficients.

Each of the models described by equations (5.1)–(5.3) were fit to the data by minimizing a cost function giving the sum of the squared difference between each of the measured $\log_{10}$-transformed $K_D$ fold-change values $y_{i,j,k}$ and their corresponding model predictions $\kappa(i,j,k)$ for all pairing-range and offset combinations *i*, *j*, and *k* with 3′-pairing lengths 4–11 bp and offset between –4 and +16 nt:

$$f_{cost,\kappa}(\boldsymbol{\alpha},\boldsymbol{\beta}) = \sum_{i=9}^{n_m-3} \sum_{j=i+3}^{n_m} \sum_{k=-4}^{+16} \left(y_{i,j,k} - \kappa(i,j,k)\right)^2, \tag{6}$$

where $\boldsymbol{\alpha}$ and $\boldsymbol{\beta}$ represent the vector of all $\alpha_{i,j}$ and all $\beta_k$ coefficients, respectively, and $n_m$ represents the length of the miRNA. This cost function was minimized with the *optim* function in R using the L-BFGS-B method, supplying the cost function and its gradient and setting the "maxit" parameter to $1 \times 10^7$. When optimizing all three models, all $\boldsymbol{\alpha}$, $\boldsymbol{\beta}$, and $\gamma$ parameters were initialized at 1.0 and bounded between 0.0 and 10.0 during the optimization.

Because the multiplicative model $\kappa_m(i,j,k)$ ($r^2$ = 0.92, 0.86, and 0.96 for let-7a, miR-1, and miR-155, respectively, Figure S4D) performed significantly better than the additive model $\kappa_a(i,j,k)$ ($r^2$ = 0.81, 0.81, and 0.94), and because the multiplicative-plus-constant model $\kappa_{mc}(i,j,k)$ provided only marginally increased performance ($r^2$ = 0.93, 0.86, and 0.96) while decreasing model interpretability (as the constant term corresponded to a benefit to binding irrespective of the manner of the pairing to the miRNA 3′ end), we selected the multiplicative model. For the purposes of interpretation of the model coefficients, we re-scaled the coefficients as follows:

$$\boldsymbol{\alpha}' = \boldsymbol{\alpha} \times \max \boldsymbol{\beta} \tag{7.1}$$

$$\boldsymbol{\beta}' = \frac{\boldsymbol{\beta}}{\max \boldsymbol{\beta}}, \tag{7.2}$$

which, because none of the coefficients were negative, caused each offset coefficient $\beta'_k$ to fall between 0.0 and 1.0, thereby corresponding to a different fractional reduction in binding energy for each offset *k*. Each re-scaled pairing range coefficient $\alpha'_{i,j}$ therefore also represented the maximum $K_D$ fold change that could be obtained by contiguous pairing to nucleotides *i* through *j*.

We estimated the model error by calculating the asymptotic covariance matrix $\boldsymbol{V}\langle\boldsymbol{\theta}\rangle$ using the standard approximation

$$\boldsymbol{V}\langle\hat{\boldsymbol{\theta}}\rangle = \frac{f_{cost,\kappa}(\hat{\boldsymbol{\theta}})}{n-p}\left(\boldsymbol{A}\langle\hat{\boldsymbol{\theta}}\rangle^T \boldsymbol{A}\langle\hat{\boldsymbol{\theta}}\rangle\right)^{-1}, \tag{8}$$

where $\hat{\boldsymbol{\theta}}$ is the vector of all optimal pairing and offset coefficients, *n* is the total number of data points, *p* is the total number of model parameters (i.e., the length of vector $\hat{\boldsymbol{\theta}}$), and $\boldsymbol{A}\langle\hat{\boldsymbol{\theta}}\rangle$

represents the matrix of partial derivatives $\frac{\partial \kappa}{\partial \theta}$ (Alper and Gelb, 1990). From the covariance matrix, the 95% confidence intervals for each model coefficient are given by

$$\widehat{\boldsymbol{\theta}} \pm t_{\alpha=0.975,\nu=n-p}\sqrt{\mathrm{diag}\big(\boldsymbol{V}\langle\widehat{\boldsymbol{\theta}}\rangle\big)}\,, \tag{9}$$

where $t_{\alpha=0.975,\nu=n-p}$ represents the *t* statistic for 97.5% confidence with *n* – *p* degrees of freedom. Because the form of the model described allows the cost function to be minimized with an infinite number of distinct solutions (where, given a particular optimal $\widehat{\boldsymbol{\theta}}$ comprised of $\widehat{\boldsymbol{\alpha}}$ and $\widehat{\boldsymbol{\beta}}$, any $\widehat{\boldsymbol{\theta}}'$ comprised of $c\widehat{\boldsymbol{\alpha}}$ and $\widehat{\boldsymbol{\beta}}/c$ is an equivalent solution), the matrix given by $\big(\boldsymbol{A}\langle\widehat{\boldsymbol{\theta}}\rangle^T\boldsymbol{A}\langle\widehat{\boldsymbol{\theta}}\rangle\big)$ is not linearly independent, and thus cannot be inverted as required in equation (8). This issue can be circumvented by arbitrarily fixing one parameter in the course of the optimization. We therefore optimized the model 21 times, fixing each $\beta_k$ coefficient at 1.0 during the optimization, determining the 95% confidence intervals for all other coefficients, and then rescaling all parameters. This led to 21 different estimates of the confidence intervals for each pairing coefficient, and 20 distinct estimates of the confidence intervals for each offset coefficient, which were averaged to produce the error estimates reported throughout the study. We note that because the parameters were re-scaled after both the optimization and confidence interval calculation, the final, re-scaled parameter values obtained were identical in each of the 21 optimization routines.

**Theoretical relationship between $K_D$ fold change and Δ*G* of 3′ pairing**

Because we were interested in the expected relationship between the measured $K_D$ fold-change values (and, by extension, the model-determined pairing coefficients), we derived their theoretical relationship by comparison of the apparent $K_D$ for a biochemical model in which only seed pairing was allowed (model A) with that in which both seed and 3′ pairing were allowed (model B). Model A is described by

$$R + A \overset{K_1}{\leftrightarrow} C_1\,, \tag{10}$$

where $R$ is the target RNA, $A$ is the unbound AGO–miRNA complex, $C_1$ is the AGO–miRNA complex seed pairing to the target RNA, and $K_1$is the equilibrium constant describing the forward reaction, defined by

$$K_1 = \frac{[C_1]}{[R][A]}\,, \tag{11}$$

where $[N]$ denotes the concentration of each species $N$. The fractional occupancy of target RNA

$R$ with model A, $\theta_A$, is given by

$$\theta_{\mathrm{A}} = \frac{[C_1]}{[R] + [C_1]}, \tag{12}$$

and when substituted for $[C_1]$ using equation (11) yields the familiar equation

$$\theta_{\mathrm{A}} = \frac{[A]}{1/K_1 + [A]}, \tag{13}$$

in which $K_1$ is the reciprocal of $K_D$.

Model B is described by

$$R + A \overset{K_1}{\leftrightarrow} C_1 \overset{K_2}{\leftrightarrow} C_2\,, \tag{14}$$

where, in addition to the terms defined for model A, $C_2$ is the AGO–miRNA complex bound to the target RNA with both seed and 3′ pairing, and $K_2$ is the equilibrium constant describing the forward reaction, defined by

$$K_2 = \frac{[C_2]}{[C_1]}\,, \tag{15}$$

which mechanistically corresponds to the AGO–miRNA complex engaging in 3′ pairing with a target to which it is already seed-paired. We note that this model does not allow formation of $C_2$ through an intermediate in which 3′ pairing occurs prior to seed pairing. Such a model was not used because 3′-autonomous pairing is considerably rarer than seed pairing and because it prevented the derivation from becoming unnecessarily complex.

The fractional occupancy of the target RNA in model B is given by

$$\theta_B = \frac{[C_1] + [C_2]}{[R] + [C_1] + [C_2]}\,, \tag{16}$$

which can be substituted for $C_2$ using equation (15) to yield

$$\theta_B = \frac{[C_1](K_2 + 1)}{[R] + [C_1](K_2 + 1)}\,, \tag{17}$$

and then further substituted for $C_1$ using equation (11) to yield

$$\theta_B = \frac{K_1[R][A](K_2 + 1)}{[R] + K_1[R][A](K_2 + 1)}\,. \tag{18}$$

This can be simplified and rearranged in an analogous form to equation (13):

$$\theta_B = \frac{[A]}{\frac{1}{K_1(K_2 + 1)} + [A]}\,, \tag{19}$$

from which it is clear that the apparent $K_D$ value is ${}^{1}/_{K_1(K_2 + 1)}$. This indicates that the expected (beneficial) fold change to the $K_D$ is $(K_2 + 1)$, and that the $K_2$, rather than the $K_D$ fold change itself, would be expected to relate to the Δ*G* through the equation $K_2 = e^{-\Delta G/RT}$. We explored incorporating this approach by fitting the model described in the prior section to $\log_{10}$[($K_D$ fold change) – 1] instead of $\log_{10}$($K_D$ fold change), but ultimately chose not to include this conversion in our analyses, because it caused many of the lowest-affinity pairing coefficients to be fit to anomalously large negative values. This occurred because those pairing coefficients were fit using $K_D$ fold-change values that in some cases closely approached 1 (i.e., the 3′ pairing provided almost no benefit), which is a regime in which small changes in measured $K_D$ fold change will lead to much larger changes in the modeled pairing coefficient. For example, a $K_D$ fold change of 1.1 would correspond to an apparent pairing coefficient of $\log_{10}(1.1 - 1) = -1$, whereas 1.01 would correspond to an apparent pairing coefficient of $\log_{10}(1.01 - 1) = -2$, or 10-fold worse binding.

**Nearest neighbor model–based prediction of 3′-compensatory pairing Δ*G***

For comparison with each pairing-range coefficient beginning at position *i* and ending at position *j*, the predicted Δ*G* of duplex formation between sequence of the miRNA beginning at position 9 and the sequence reverse-complementary to miRNA positions *i*–*j*, with no non-complementary nucleotides appended to either terminus, was calculated using RNAduplex, as part of the ViennaRNA package, through its Python interface (Lorenz et al., 2011).

**Calculation of mismatch coefficients**

We extended the thermodynamic model of 3′-end binding efficacy to include seed-mismatch effects, by using as input data the $\log_{10}$($K_D$ fold-change) values measured for each of the 18 8mer-mismatch sites separately. This model took the form of:

$$\kappa_2(i,j,k,l) = \alpha_{i,j}\beta_k\delta_l. \tag{20}$$

where $\alpha_{i,j}$ and $\beta_k$ represented the pairing and offset preferences as before, and $\delta_l$ represented the additional set of 18 seed-mismatch coefficients. The updated cost function was therefore

$$f_{cost,\kappa_2}(\boldsymbol{\alpha},\boldsymbol{\beta},\boldsymbol{\delta}) = \sum_{i=9}^{n_m-3}\sum_{j=i+3}^{n_m}\sum_{k=-4}^{+16}\sum_{l=1}^{18}\left(y_{i,j,k,l} - \kappa_2(i,j,k,l)\right)^2, \tag{21}$$

where $\boldsymbol{\delta}$ represents the vector of all $\delta_l$ coefficients. The optimization was performed identically as before, with the $\gamma$ parameters initialized at 1.0, and bounded between 0.0 and 10.0 during the optimization. After the optimization, the coefficients were re-scaled as

$$\boldsymbol{\alpha}' = \boldsymbol{\alpha} \times \max \boldsymbol{\beta} \times \text{mean}\, \boldsymbol{\delta} \tag{22.1}$$

$$\boldsymbol{\beta}' = \frac{\boldsymbol{\beta}}{\max \boldsymbol{\beta}} \tag{22.2}$$

$$\boldsymbol{\delta}' = \frac{\boldsymbol{\delta}}{\text{mean}\, \boldsymbol{\delta}}\ , \tag{22.3}$$

which preserved the same interpretation of each $\alpha'_{i,j}$ and $\beta'_k$ coefficient as for the prior model, and further parameterized each $\delta'_l$ to represent the multiplicative deviation in binding caused by each seed-mismatch type *l*, such that the average effect of all 18 mismatches was that of being multiplied by 1.

The 95% confidence intervals of this model also used equations (8) and (9). However, because this model was the product of three sets of categorical coefficients, one coefficient from each of two sets was required to be fixed while performing the error determination. We therefore optimized the model 21 × 18 times, fixing one $\beta_k$ coefficient at 1.0 and one $\delta_k$ at 1.0 during the optimization, determining the 95% confidence intervals for all other coefficients, and then rescaling all parameters. This led to 21 × 17 = 357 different estimates of the confidence intervals for each seed-mismatch coefficient, which were averaged to produce the error estimates reported throughout the study.

**Empirical assessment of the influence of seed-type on 3′ pairing with the random-library AGO-RBNS experiments**

When analyzing the effects of seed type on $K_D$ fold change in the random-library experiments, we first attempted to apply the modeling approach as used when analyzing the data from the programmed-library experiments. However, we found that the read counts assigned to each possible pairing, offset, and seed-mismatch combination were too sparse, such that modeling using these data could not be reliably performed. We therefore analyzed the differences in the benefit of 3′-supplementary pairing between each of the six canonical sites (8mer, 7mer-m8, 7mer-A1, 6mer, 6mer-m8, and 6mer-A1) as well as a representative 3′-compensatory site given by summing the read counts of all 18 8mer-mismatch sites.

To compare these sites, we first took, for each miRNA, all 3′ pairing possibilities with lengths of 4 or 5 bp and pairing to miRNA positions 9–18, and determined for each of these possibilities the offset that imparted the maximum average $\log_{10}$($K_D$ fold change), thereby constructing a 7 (site types, i.e., the six canonical sites and a combination of the 18 8mer-mismatch sites) × 20 (pairing range) matrix of $\log_{10}$($K_D$ fold change) values. This matrix was then sorted by the average $\log_{10}$($K_D$ fold change) of each pairing range, and then this value was subtracted from each column, such that the values within each column reported on the deviation of each site type from the average, for that pairing-and-offset possibility. These deviations were then averaged for each of the seven site types over the top five (i.e., the top quartile) of pairing-and-offset possibilities, to give the empirical contribution of each site type to 3′-binding affinity, in comparison to that of the average. These values are plotted for all six miRNAs in Figure 4H, and the data tables from which they were calculated are visualized in Figure S11, with the columns used as input for the final averaging indicated.

**Assignment of miRNA sites with imperfect 3′ pairing within reads from the programmed libraries**

To calculate the effect of a mismatch, bulge, or deletion on the binding affinity of a 3′ site, the site counting was repeated for each fully paired 3′ site while also counting each of its derivatives containing either a single mismatched, bulged, or deleted nucleotide (hereafter referred to as "imperfect 3′ sites"). This was done to reduce the total number of sites being simultaneously counted and used to calculate relative $K_D$ values, and as well to reduce assignment problems owing to any imperfect 3′ site from one region of the miRNA being sequence-identical to that from any other region of the miRNA.

The site counting was performed similarly to that of the fully paired 3′ sites, with some differences: those reads containing a seed site at the programmed region were still assigned to one of four categories, with the definition of a 3′ site expanded to include any fully paired 3′ site of length 4–11 nt pairing to the miRNA 3′ end in addition to any imperfect derivatives of that site. The categorization of each read proceeded through the following steps: 1) Any seed sites were identified, looking within the entire random-programmed-random region with 3 nt of constant sequence appended to the 5′ and 3′ ends of the read. 2) The 28-nt segment comprising 3 nt of the 5′-constant sequence and 25 nt of random sequence was queried for any instances of any imperfect 3′ sites, with any deleted- or mismatched-nucleotide sites that were contained with another mismatched- or bulged-nucleotide site not counted (e.g., the let-7a 11mer-m11–20 with a mismatched U at position 20 is inherently contained within the 11-mer-m11–20 with a bulged U between positions 19 and 20, but only the bulged-nucleotide version of the site would be recorded). Any imperfect 3′ sites were also queried to make sure that the nucleotides on each side of the site were not complementary to the next corresponding positions of the miRNA guide. We note that if such an imperfect 3′ site was found that failed these criteria, any identified fully paired 3′ sites were not counted toward that read. 3) The 28-nt segment comprising 3 nt of the 5′-constant sequence and 25 nt of random sequence was queried for the longest fully paired 3′ site(s), retaining multiple 3′ sites if more than one was of the longest length. If at least one fully paired 3′ site >4 nt in length and not contained within a seed site (if such sites were present) was identified, and if at least one imperfect 3′ sites was identified, the length of the fully paired 3′ site(s) was compared to that of the imperfect 3′ site(s), and the 3′ site(s) that was longer was retained. Lastly, if the fully paired 3′ site(s) was ≥11 nt in length, neither the fully paired nor imperfect 3′ site(s) were counted. 4.) The read was then assigned one of the four categories, and the read count split as described when assigning contiguous 3′ sites.

The relative $K_D$ values used in Figures 7 and S12 were calculated using the summed counts of each site over all 18 mismatch sites in the programmed region. In order to mitigate noise in the measurement of affinities for individual imperfect 3′ sites, the $K_D$ fold-change values of each fully paired site and each of its imperfect sites at particular offset values were substituted with the geometric mean of the $K_D$ fold change of those sites over the three contiguous offset values centered on that offset value. The offset value used for reporting of the imperfect $K_D$ fold-change values was the offset value between –2 and +6 nt for which the $K_D$ fold-change of the fully paired site was the greatest.

When performing the positional mismatch analyses of Figure 7E, $\Delta\Delta G$ values corresponding to sites in which the position of a particular bulged nucleotide was ambiguous were used to calculate the average $\Delta\Delta G$ value for each of the inter-nucleotide positions at which it might occur. Conversely, these sites were omitted from comparisons of terminal bulges and mismatches of Figure 7F.

**Assignment of miRNA sites with imperfect 3′ pairing within reads from the random-sequence libraries**

When counting imperfect 3′ sites within the random-library reads, we similarly preformed the read counting and relative $K_D$ fitting for each fully paired site and its imperfect derivatives. Individual reads were queried for all seed sites and fully paired 3′ sites as described for the fully paired site counting with the random-library experiments, in addition to being queried for any imperfect 3′ sites derived from a fully paired site. If a read contained at least one seed site and at least one fully paired or imperfect 3′ site, each 3′ site was checked for any amount of overlap with any seed sites. If a fully paired 3′ site overlapped a seed site with the 3′ site being the 5′-most site, the 3′ site was trimmed to exclude the region overlapping the seed site, and retained if the site was still ≥4 nt in length. As before, any fully paired 3′ sites that either overlapped any seed sites from the 3′ end, contained a seed site within them, or were entirely contained within a seed site, were discarded. Any imperfect 3′ sites identified were examined to ensure that read positions just outside their limits were not complementary to the corresponding miRNA positions, and as well that they did not overlap any of the seed sites within the read.

If one or more 3′ site (either fully paired or imperfect) remained, all of the fully paired 3′ sites of length equal to that of the longest fully paired 3′ site in the read were retained. If at least one fully paired and at least one imperfect 3′ site remained, the length of the fully paired 3′ site(s) was compared to that of the fully paired site from which the imperfect site(s) was derived, and if it was shorter, the imperfect 3′ site(s) was retained, and the fully paired site(s) was discarded. If the fully paired 3′ site(s) was longer than the fully paired site from which the imperfect site(s) was derived, the fully paired site(s) was retained, and the imperfect site(s) was discarded. All possible bipartite sites associated with that read were then enumerated, whereby each seed site was considered to form a bipartite site with each 3′ sites that was fully 5′ of that seed site. The read was then split among any bipartite sites identified. In the event that no bipartite sites were identified, the read was split among all the seed and 3′ sites equally. While this procedure ensured equal partitioning of read counts in the case of multiple seed or 3′ sites within a read, in practice only a small fraction of reads contained either multiple seed sites and a 3′ site or a seed site and multiple 3′ sites.

The $K_D$ fold-change values associated with each imperfect site were calculated from the ratio of the relative $K_D$ value of the site when summing the counts for all 18 8mer-mismatch sites to the geometric mean of the relative $K_D$ values of each of the 18 8mer mismatch sites calculated individually. As with the relative $K_D$ values corresponding to the imperfect sites derived from the programmed libraries, the values reported for each of the perfect and imperfect sites reported in Figure S13 were the geometric mean of the values from the three contiguous offsets at which the $K_D$ fold change of the fully paired site was the greatest.

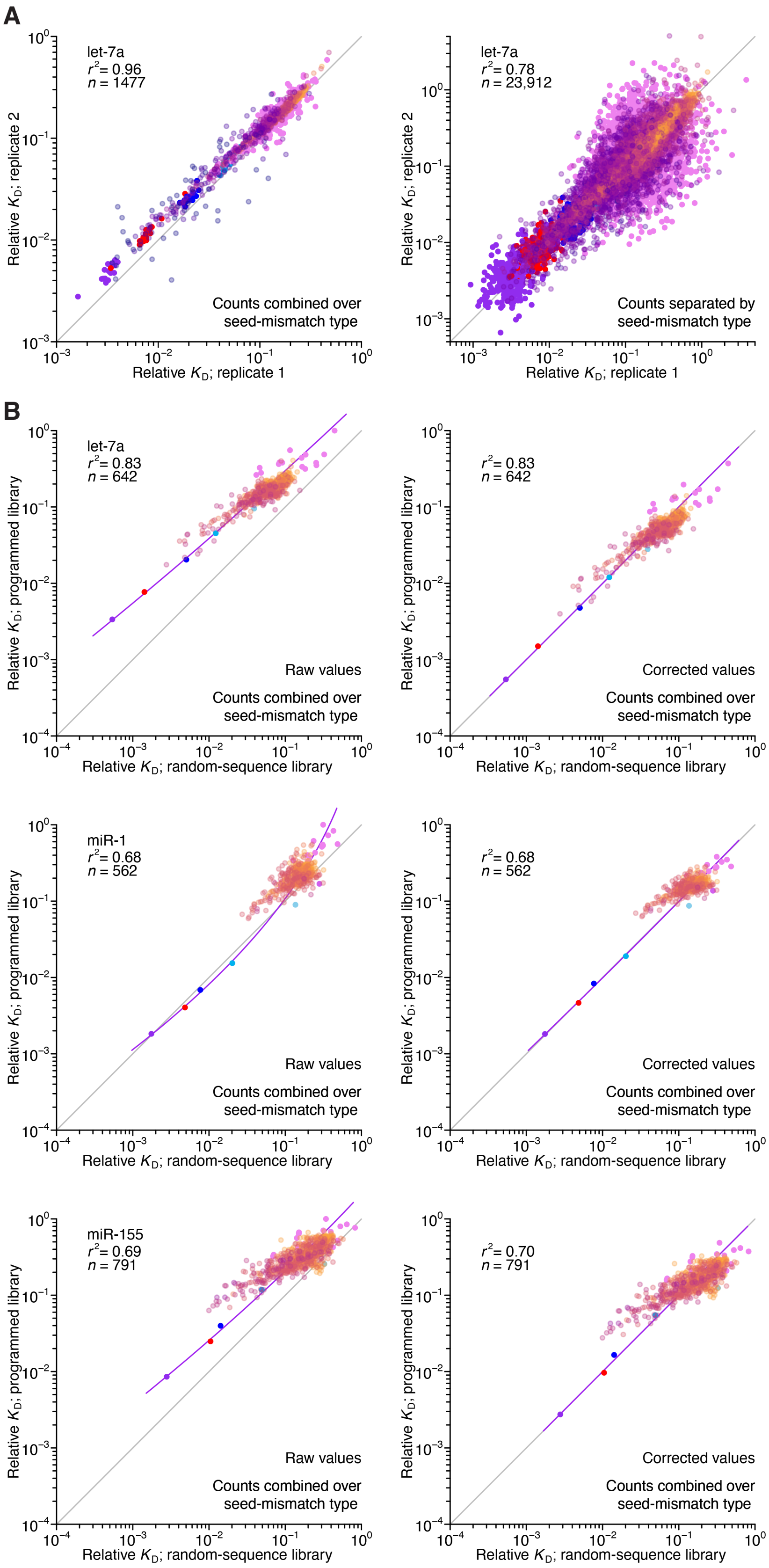

A
let-7a
r² = 0.96
n = 1477
Relative KD; replicate 2
Relative KD; replicate 1
Counts combined over seed-mismatch type
let-7a
r² = 0.78
n = 23,912
Counts separated by seed-mismatch type
B
let-7a
r² = 0.83
n = 642
Relative KD; programmed library
Relative KD; random-sequence library
Raw values
Counts combined over seed-mismatch type
r² = 0.83
n = 642
Corrected values
miR-1
r² = 0.68
n = 562
r² = 0.68
n = 562
miR-155
r² = 0.69
n = 791
r² = 0.70
n = 791

**Figure S1. Reproducibility of AGO-RBNS with programmed libraries and correspondence with random-sequence libraries**

(**A**) Pairwise comparison of replicate relative $K_D$ values measured using the let-7-programmed library and AGO2–let-7a when combining (left) or separating (right) reads based on the identity of the seed-mismatch site at the programmed region of the library. Each of the two replicate experiments was performed with independent preparations of both the library and the purified AGO–miRNA complex. The relative $K_D$ values correspond to both seed sites and 3′-compensatory sites spanning 4 (orange) to 11 (dark blue) contiguous base pairs in length. The $r^2$ value reports on the coefficient of determination between the log-transformed relative $K_D$ values of each replicate.

(**B**) Pairwise comparison of the relative $K_D$ values measured for let-7a (top), miR-1 (middle), and miR-155 (bottom) in the programmed-library experiments to that of the random-library experiments, prior to (left) and after (right) correction of each of the programmed library–derived measurements according to those of the random-library experiments using LOESS. Otherwise, this panel is as in (A).

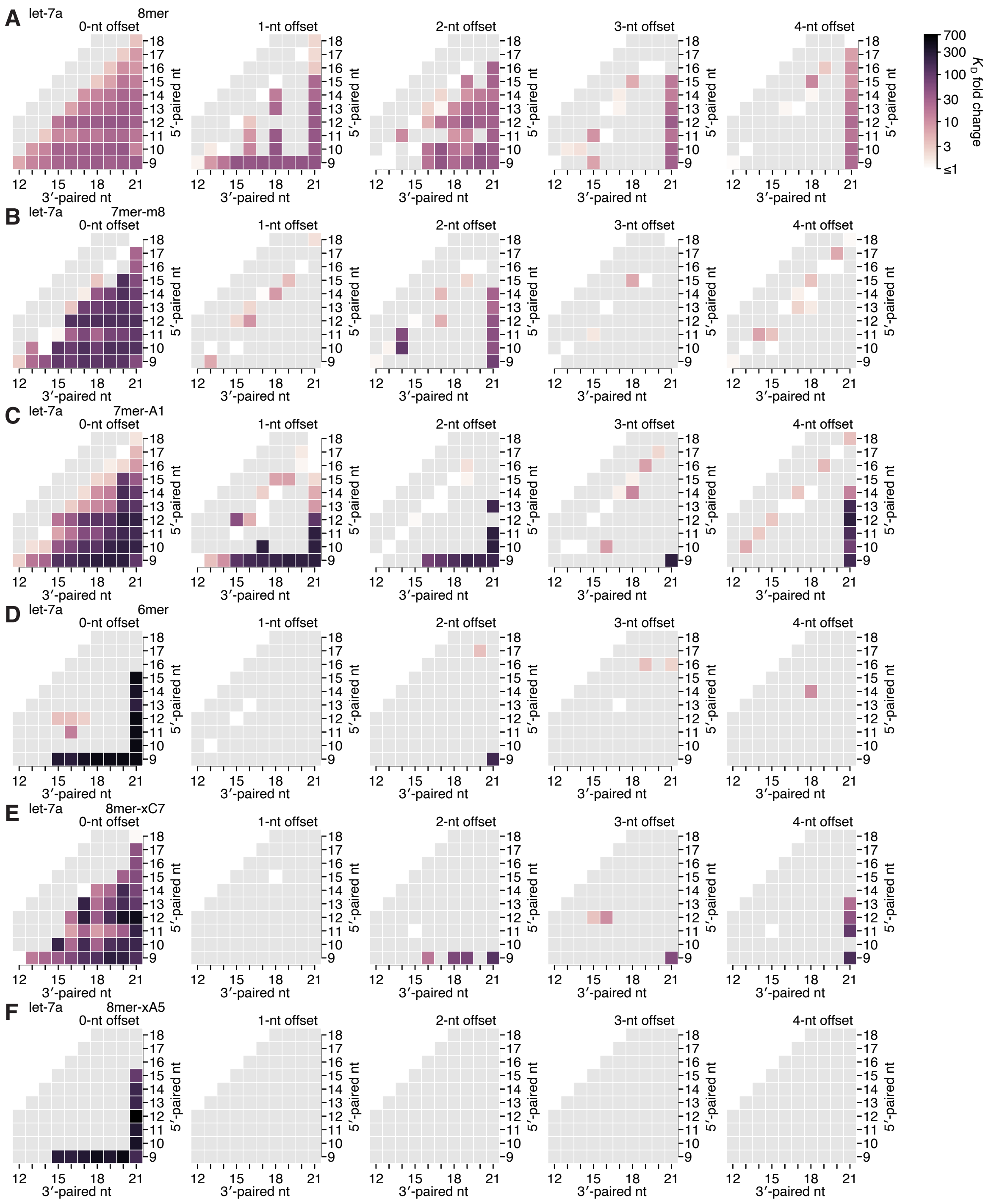


**Figure S2. Re-analysis of binding experiments performed in Becker et al. (2019)**

(**A**–**F**) Partial affinity profiles of the let-7a 3′ region in the context of the 8mer (A), 7mer-m8 (B), 7mer-A1 (C), 6mer (D), 8mer-xC7 (E), and 8mer-xA5 (F) seed sites, using binding affinity measurements calculated using imaging-based, high-throughput single-molecule experiments (Becker et al., 2019). Each cell indicates the fold change in $K_D$ attributed to a 3′ site with indicated length, position, and offset of pairing. In (A–E), the fold change is with respect to the geometric mean of the measured $K_D$ values for all target RNAs with the corresponding seed site and <4 bp of 3′ pairing, and in (F), the fold change is with respect to 10 nM, because there were no target RNAs with an 8mer-xA5 site and no 3′ pairing. Gray boxes indicate pairing possibilities for which there were no data. Otherwise, this panel is as in Figure 2E.

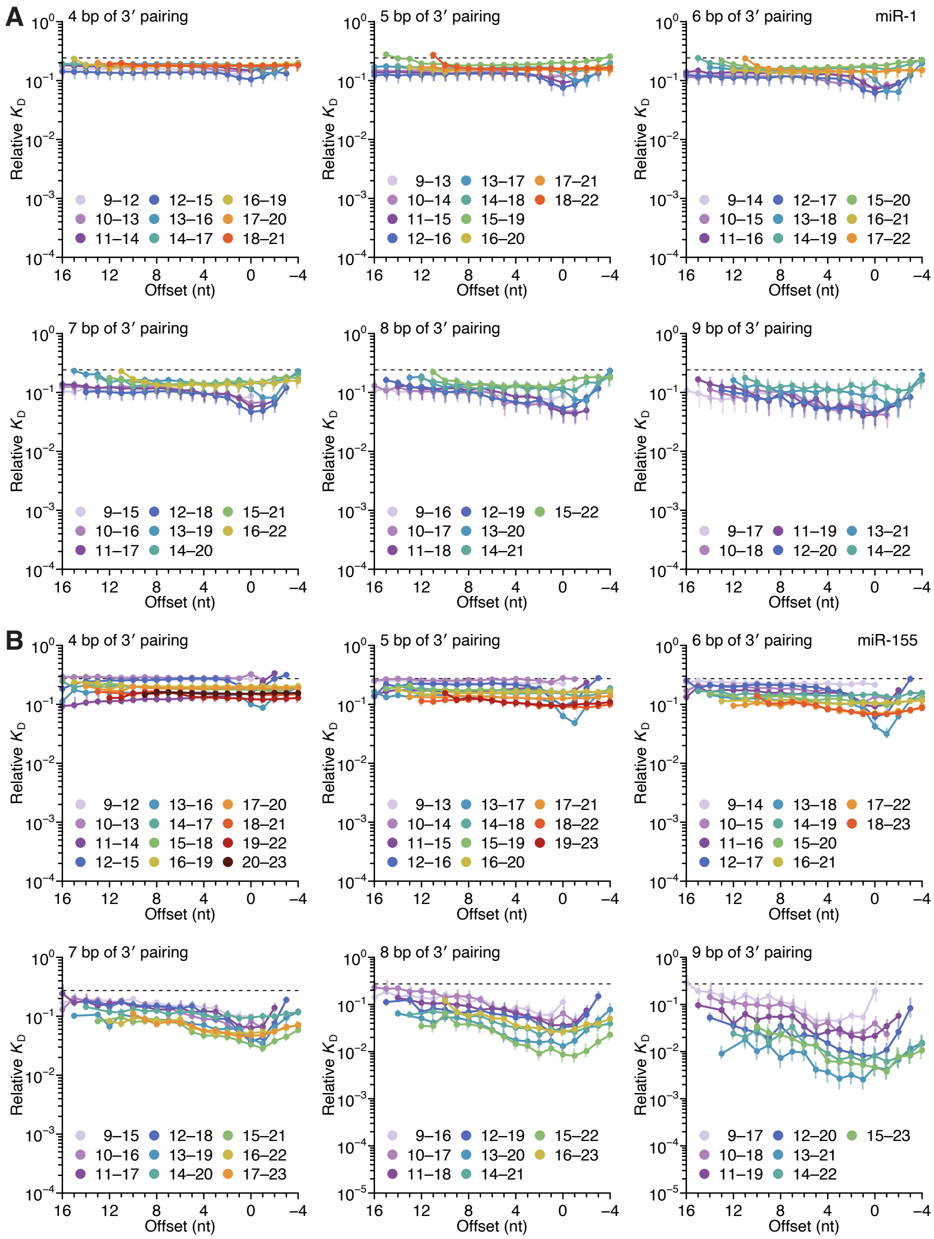


**Figure S3. Length, position, and offset trends of miR-1 and miR-155 indicate one binding mode**

(**A** and **B**) The dependency of miR-1 (A) and miR-155 (B) 3′ pairing on pairing length, position, and offset. Each panel shows the relative $K_D$ values for 3′ pairing of a specified length over a range of positions and offsets, spanning positions 9 (light violet) to 18 (red) wherever possible. Otherwise, this panel is as in Figure 2C.

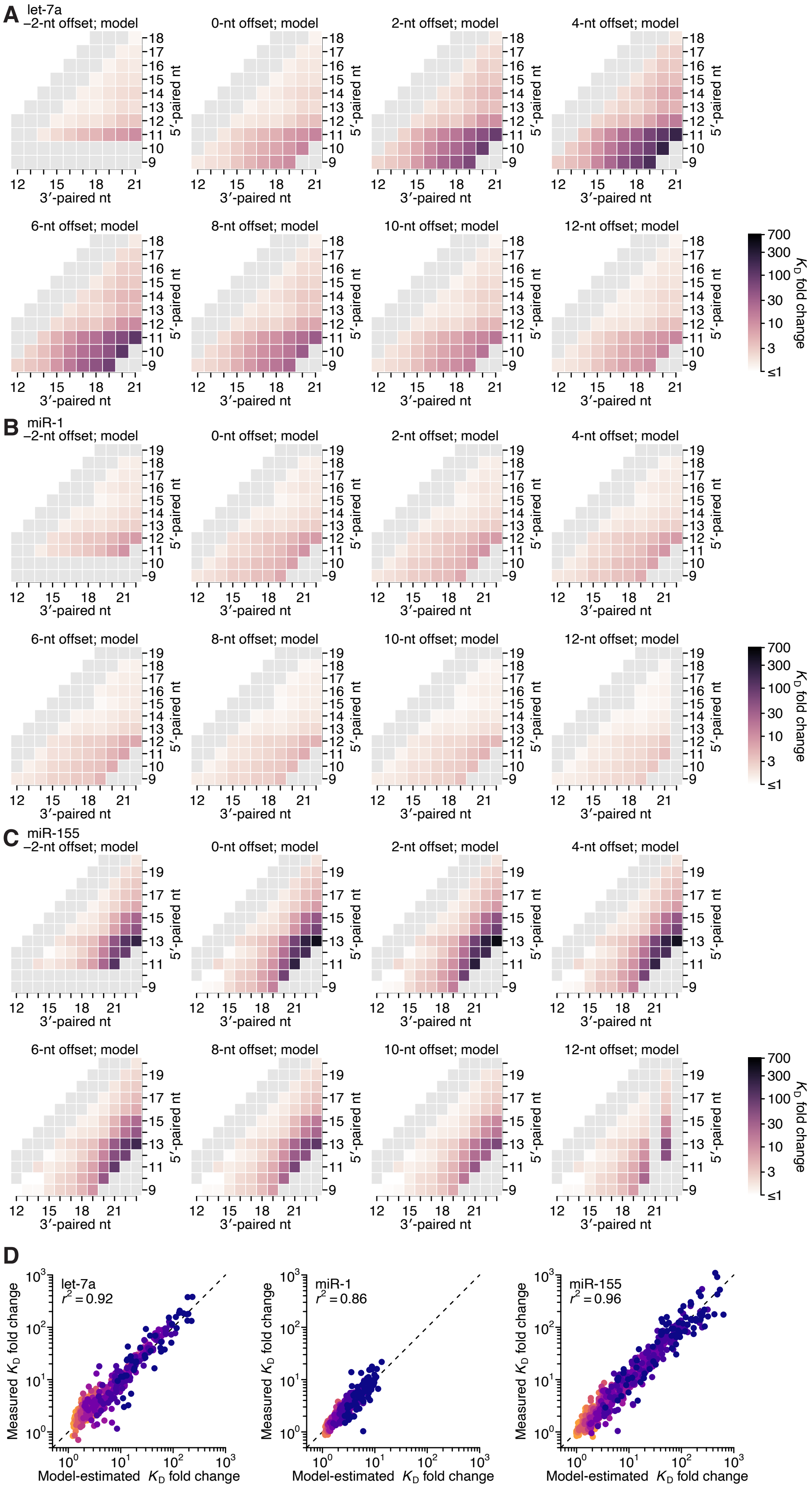
A
let-7a
−2-nt offset; model
0-nt offset; model
2-nt offset; model
4-nt offset; model
6-nt offset; model
8-nt offset; model
10-nt offset; model
12-nt offset; model
5′-paired nt
3′-paired nt
B
miR-1
C
miR-155
700
300
100
30
10
3
≤1
$K_D$ fold change
D
let-7a
$r^2$ = 0.92
miR-1
$r^2$ = 0.86
miR-155
$r^2$ = 0.96
Measured $K_D$ fold change
Model-estimated $K_D$ fold change

**Figure S4. Model prediction of 3′-compensatory pairing**

(**A**–**C**) Model-predicted affinity profiles of the let-7a (A), miR-1 (B), and miR-155 (C) 3′ regions. Gray cells correspond either to pairing lengths <4 bp or >11 bp, or to pairing ranges for which there were no measured $K_D$ fold-change values for comparison. Otherwise, these panels are as in Figure 2E.

(**D**) Pairwise comparison of the model-predicted and measured $K_D$ fold-change values for let-7a (left), miR-1 (middle), and miR-155 (right), with sites that have 4 (orange) to 11 (dark blue) contiguous bp of 3′ pairing. The $r^2$ reports on the coefficient of determination between the log-transformed predicted and measured values.

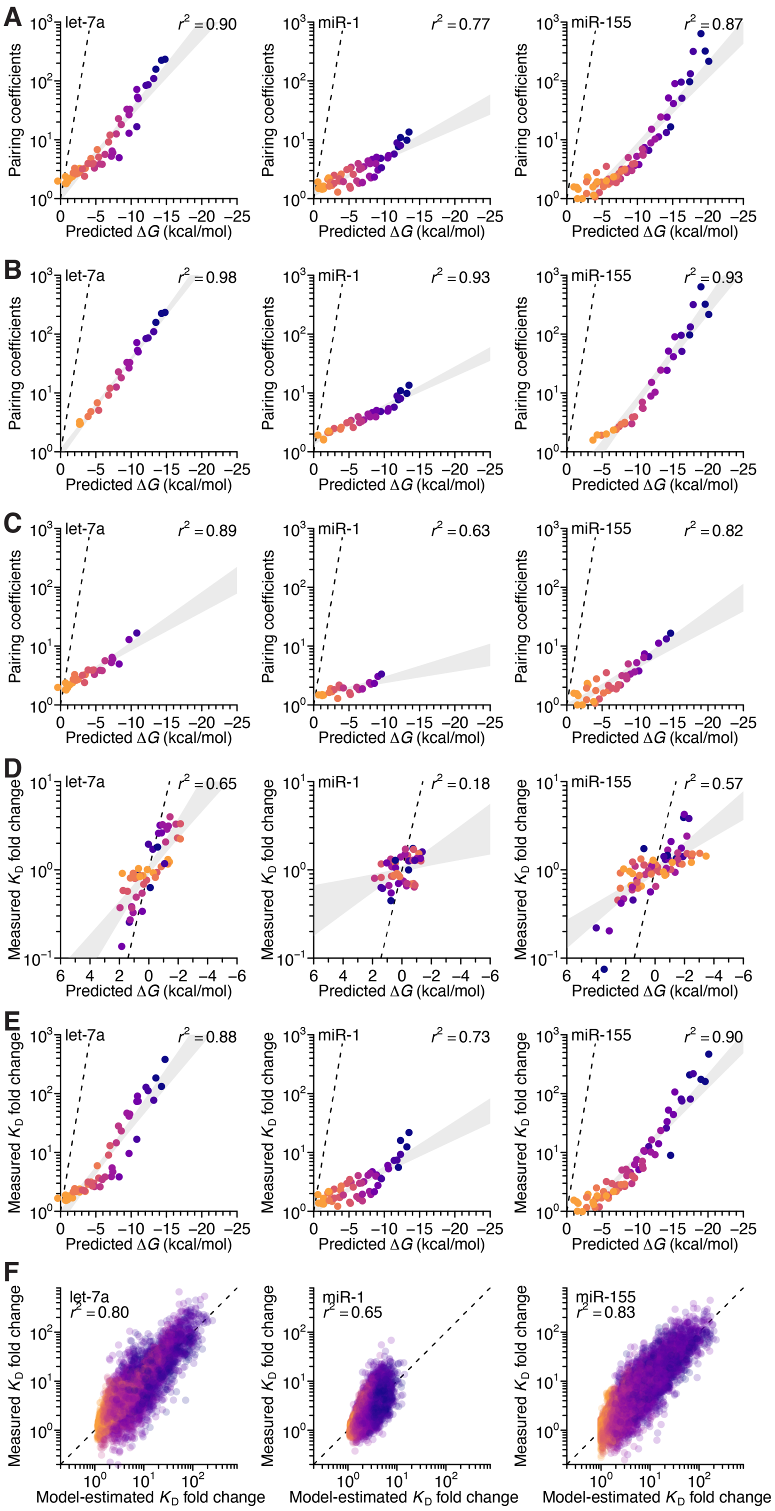


**Figure S5. Analysis of the correspondence of pairing coefficients with predicted Δ*G*, and performance of seed mismatch–effect model**

(**A**) The relationship between the model-derived pairing coefficients (Figures 4A–4C, left) and the predicted Δ*G* values (Figure 4D), when not controlling for length. Otherwise, this panel is as in Figure 4E.

(**B** and **C**) The relationship between the model-derived pairing coefficients (Figures 4A–4C, left) and the predicted Δ*G* values (Figure 4D), when separating the pairing based on whether it includes (B) or excludes (C) pairing to nucleotide 11 for let-7a (left), 12 for miR-1 (middle), and 20 for miR-155 (right). Otherwise, these panels are as in (A).

(**D**) The relationship between $K_D$ fold-change values measured at the optimal offset for each miRNA and their corresponding predicted Δ*G* values (Figure 4D), when controlling for length. Otherwise, this panel is as in Figure 4E.

(**E**) The relationship between $K_D$ fold-change values measured at the optimal offset for each miRNA and their corresponding predicted Δ*G* values (Figure 4D), when not controlling for length. Otherwise, this panel is as in (A).

(**F**) Pairwise comparison of the model-predicted and measured $K_D$ fold-change values for let-7a (left), miR-1 (middle), and miR-155 (right), when using an expanded model with pairing, offset, and seed-mismatch coefficients. Points are colored according to the length of their 3′ pairing, which ranged from 4 (orange) to 11 (dark blue) contiguous bp. The $r^2$ reports on the coefficient of determination between the log-transformed predicted and measured values.

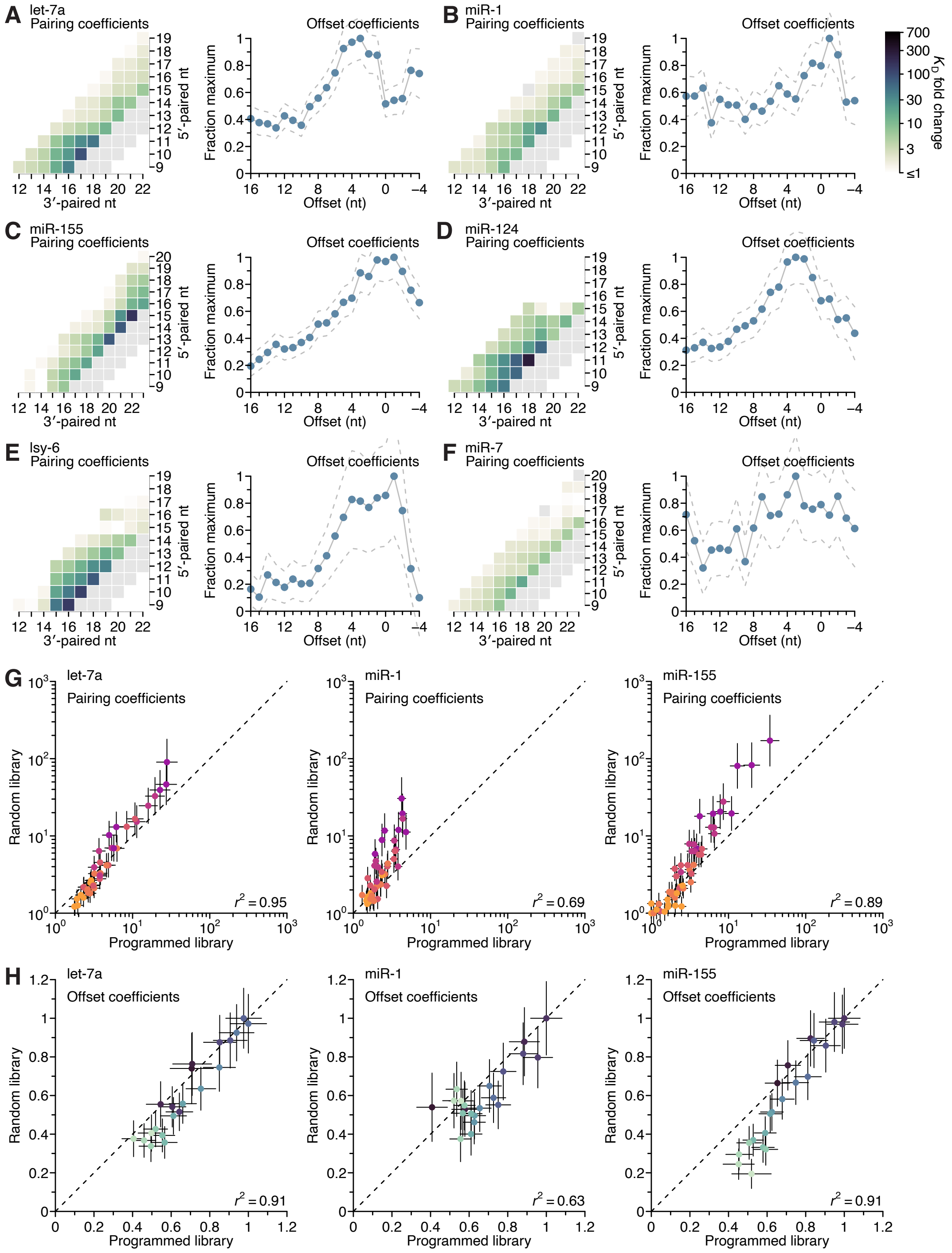


**Figure S6. Distinct pairing and offset preferences of different miRNAs in random-sequence AGO-RBNS experiments**

(**A**–**F**) Model-based analyses of 3′-pairing preferences of let-7a (A), miR-1 (B), miR-155 (C), miR-124 (D), lsy-6 (E), and miR-7 (F), for sites 4–8 nt in length, using data from previously reported, random-sequence AGO-RBNS experiments. Otherwise, these panels are as in Figure 4A, left and middle-left.

(**G** and **H**) Pairwise comparison of the pairing (G) and offset (H) coefficients derived from the programmed and random-sequence libraries for let-7a (left), miR-1 (middle), and miR-155 (right), for 3′ pairing of lengths 4–11 bp. The $r^2$ reports on the coefficient of determination between the log-transformed values for each comparison.

Inspection of pairing preferences of let-7a, miR-1, and miR-155, as indicated by their pairing and offset coefficients derived from the random sequence–library data, revealed distinguishing features consistent with those derived from the programmed-library data, including: the importance of pairing to position 11 of let-7a and position 12 of miR-1, the increased optimal offset of let-7a, and the relative ordering of the maximal benefit of 3′ pairing, with that of miR-155 exceeding that of let-7a, which exceeded that of miR-1 (Figures S6A–S6C).

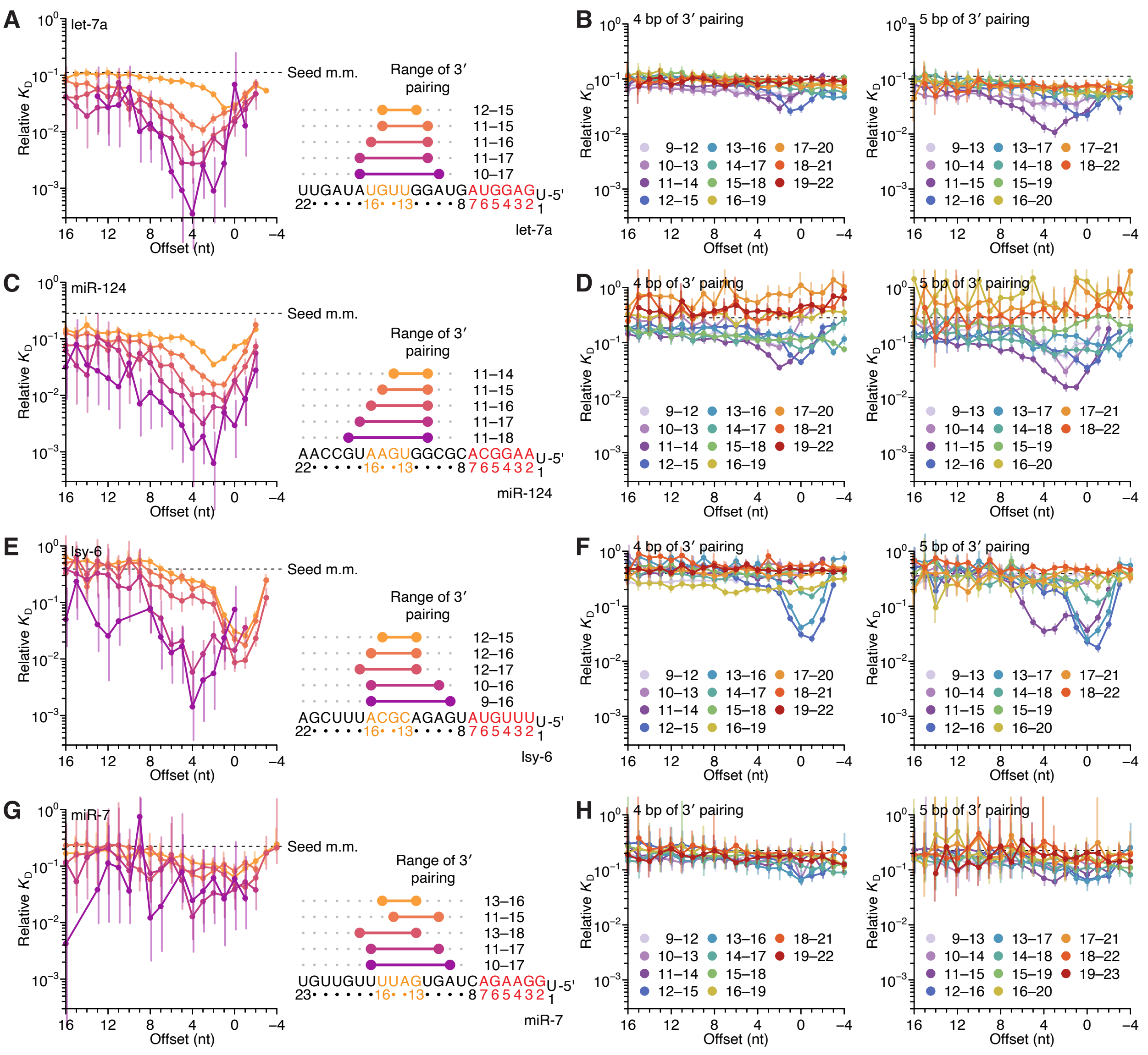


**Figure S7. Identification of two binding modes for most miRNAs in random-sequence AGO-RBNS experiments**

(**A**) Relative $K_D$ values of let-7a 3′-compensatory sites that had optimally positioned 3′ pairing of lengths 4–8 bp, as measured with data obtained previously from fully randomized libraries (McGeary et al., 2019). Otherwise, this panel is as in Figure 2B.

(**B**) The dependency of let-7a 3′ pairing affinity on pairing length, position, and offset, for 3′ pairing of 4 (left) and 5 (right) bp of pairing. Otherwise, this panel is as in Figure 2C.

(**C–H**) Equivalent analyses to those of (A) and (B), performed with miR-124 (C and D), lsy-6 (E and F), and miR-7 (G and H), as measured with data obtained previously from fully randomized libraries.

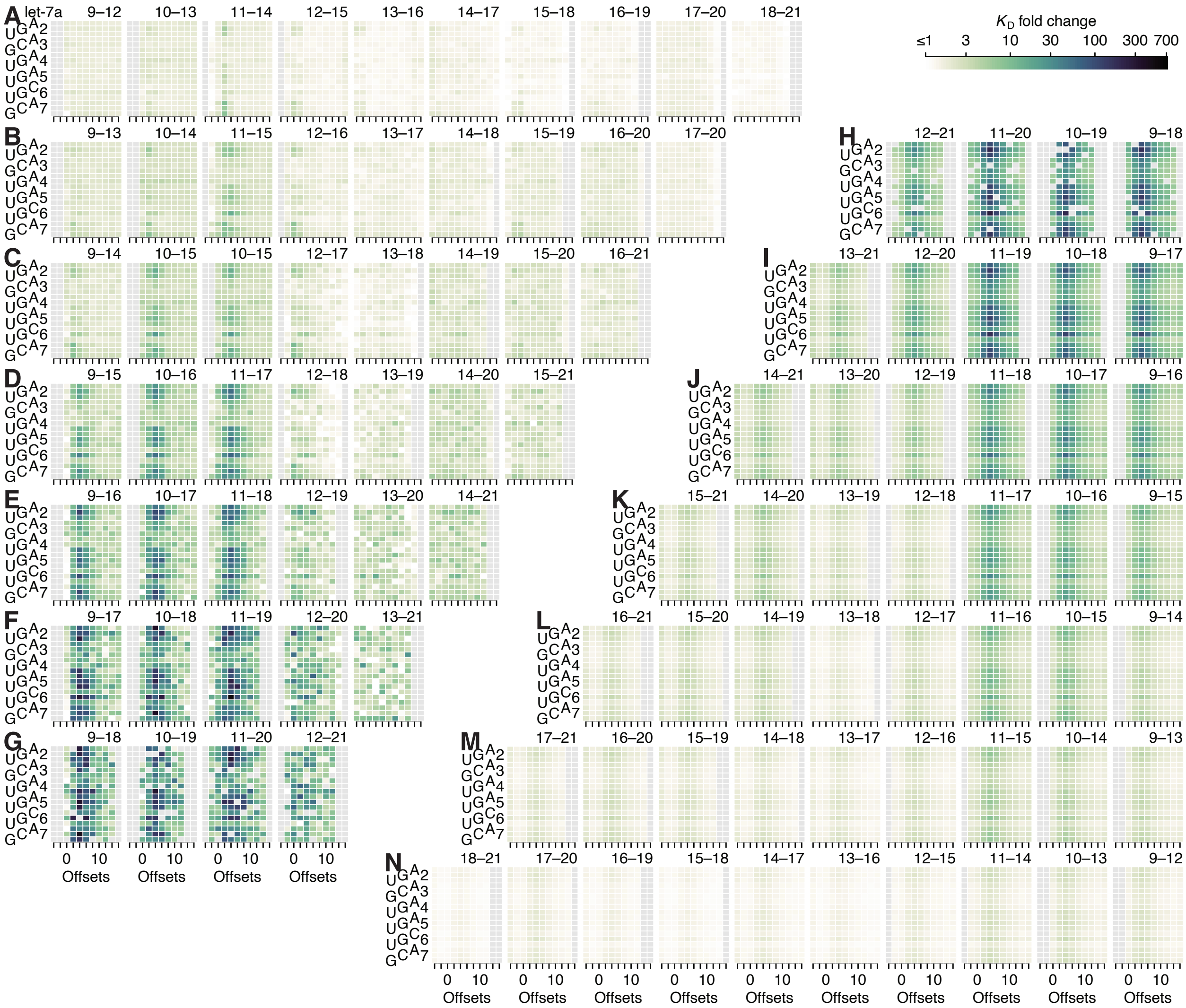

**Figure S8. Model prediction of pairing, offset, and seed-mismatch effects for let-7a**

(**A–N**) Visual representation of the model performance evaluated in Figure S5F, left. The heat maps within the upper-left triangle (A–G) show the measured $K_D$ fold-change values, while the heat maps within the lower-right triangle (H–N) show the model predictions. The two triangular arrays of heat maps are rotationally symmetric. Each individual heat map describes all of the variation associated with one defined stretch of 3′ pairing. Within each heat map, each row corresponds to a different seed mismatch type, with the left-hand numbers referring to the seed-mismatch position of every three rows, the staggered left-hand letters designating the mismatch identity of each row, and the columns each corresponding to a different even-valued offset of pairing between −4 and +16 nt.

The seed-mismatch coefficients were modeled to influence the affinity of 3′ pairing as a function of the amount of affinity associated with each 3′ pairing architecture, which varied between miRNAs. Thus, the range of 0.50, 0.48, and 0.57 observed for seed-mismatch coefficients for let-7a, miR-1, and miR-155, respectively (Figure 4F), corresponded to 9.2-, 2.6-, and 11.2-fold predicted variation in $K_D$ fold change, respectively, for each miRNA in the context of its most favorable pairing and offset, with the lower impact on miR-1 attributed to the lower amount of affinity attainable by its 3′ pairing. These predicted effects generally agreed with those observed when examining affinities of the top quartile of 3′ sites in the context of their optimal offset, which respectively varied by 14.9-, 5.1-, and 6.5-fold depending on seed-mismatch identity. Furthermore, visual inspection of the trends in observed 3′-site affinities confirmed the increased effect of seed mismatches for higher-affinity 3′ sites (Figures S8–S10). For example, only a few of the 4-nt 3′ sites to let-7a were sensitive to the particular seed-mismatch type (Figure S8A), whereas for 8-nt sites, more positions and offsets exhibited such variation, and these were positions and offsets with higher average affinities (Figure S8E).

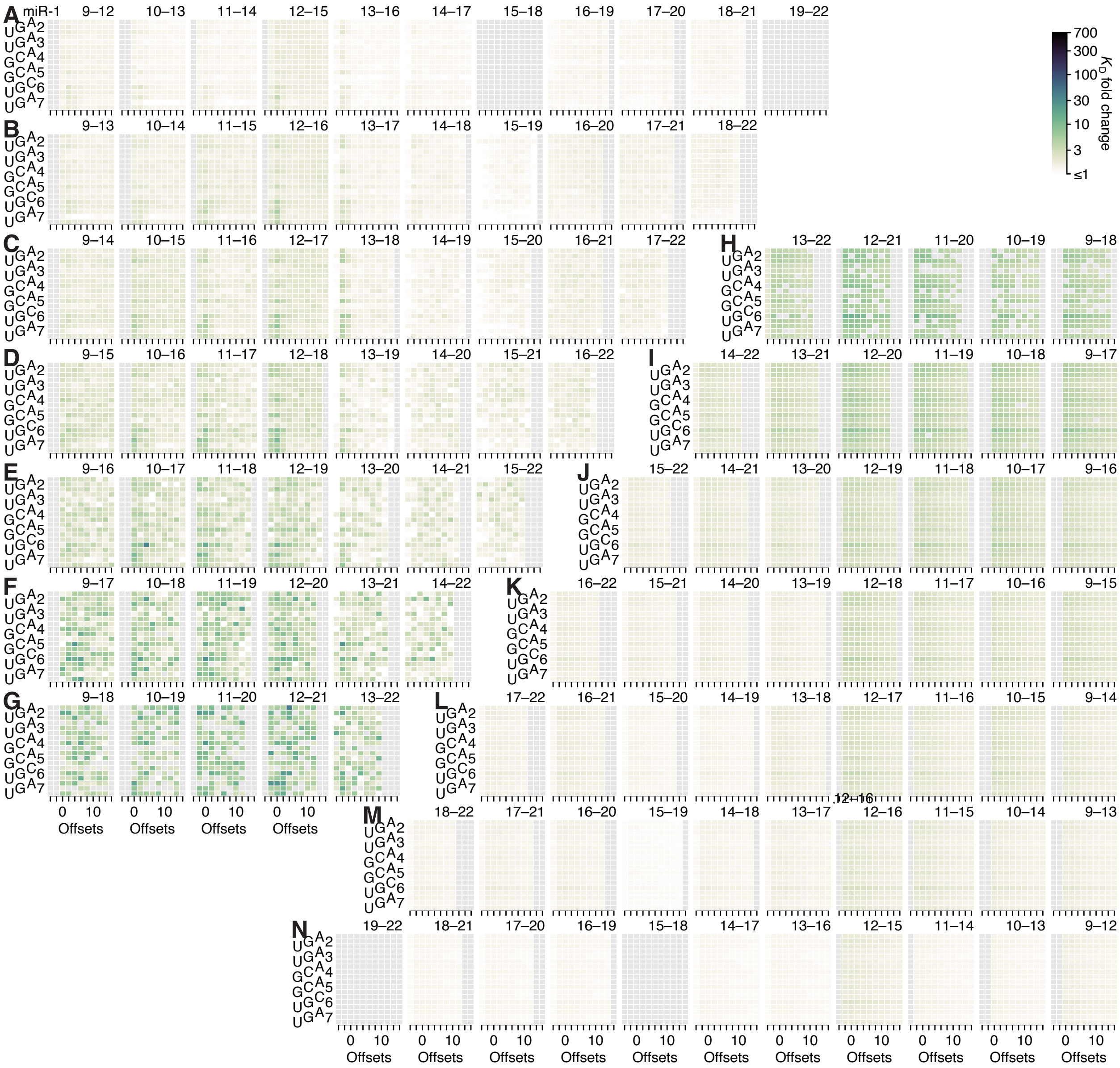

**Figure S9. Model prediction of pairing, offset, and seed-mismatch effects for miR-1**

(**A**–**N**) Visual representation of the model performance evaluated in Figure S5F, middle. No model-predicted or measured $K_D$ fold-change values are reported for pairing to positions 15–18 or 19–22 because both segments have the same sequence and are thus indistinguishable. Otherwise, this figure is as in Figure S8.

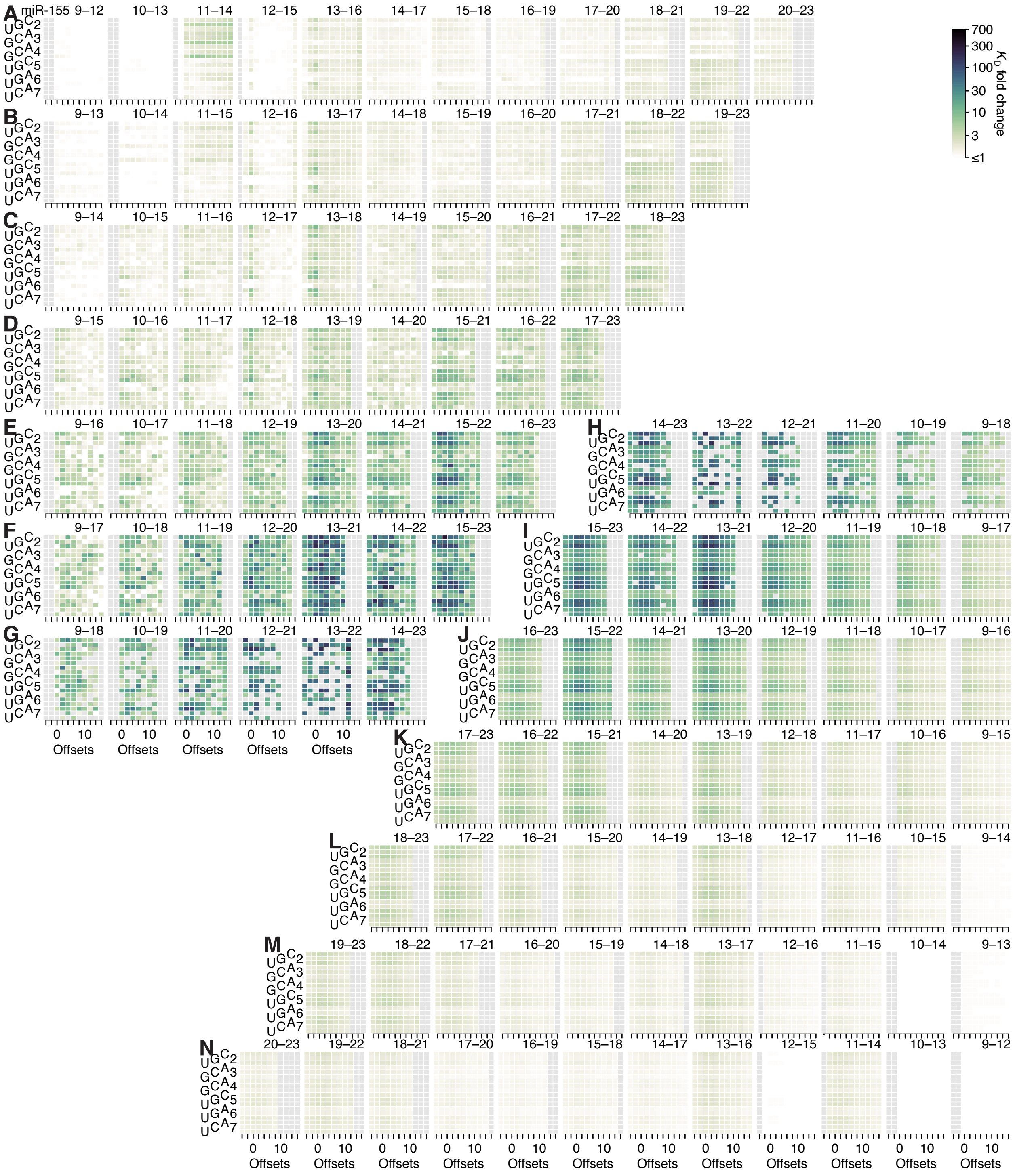

**Figure S10. Model prediction of pairing, offset, and seed-mismatch effects for miR-155**

(**A**–**N**) Visual representation of the model performance evaluated in Figure S5F, right. Otherwise, this figure is as in Figure S8.

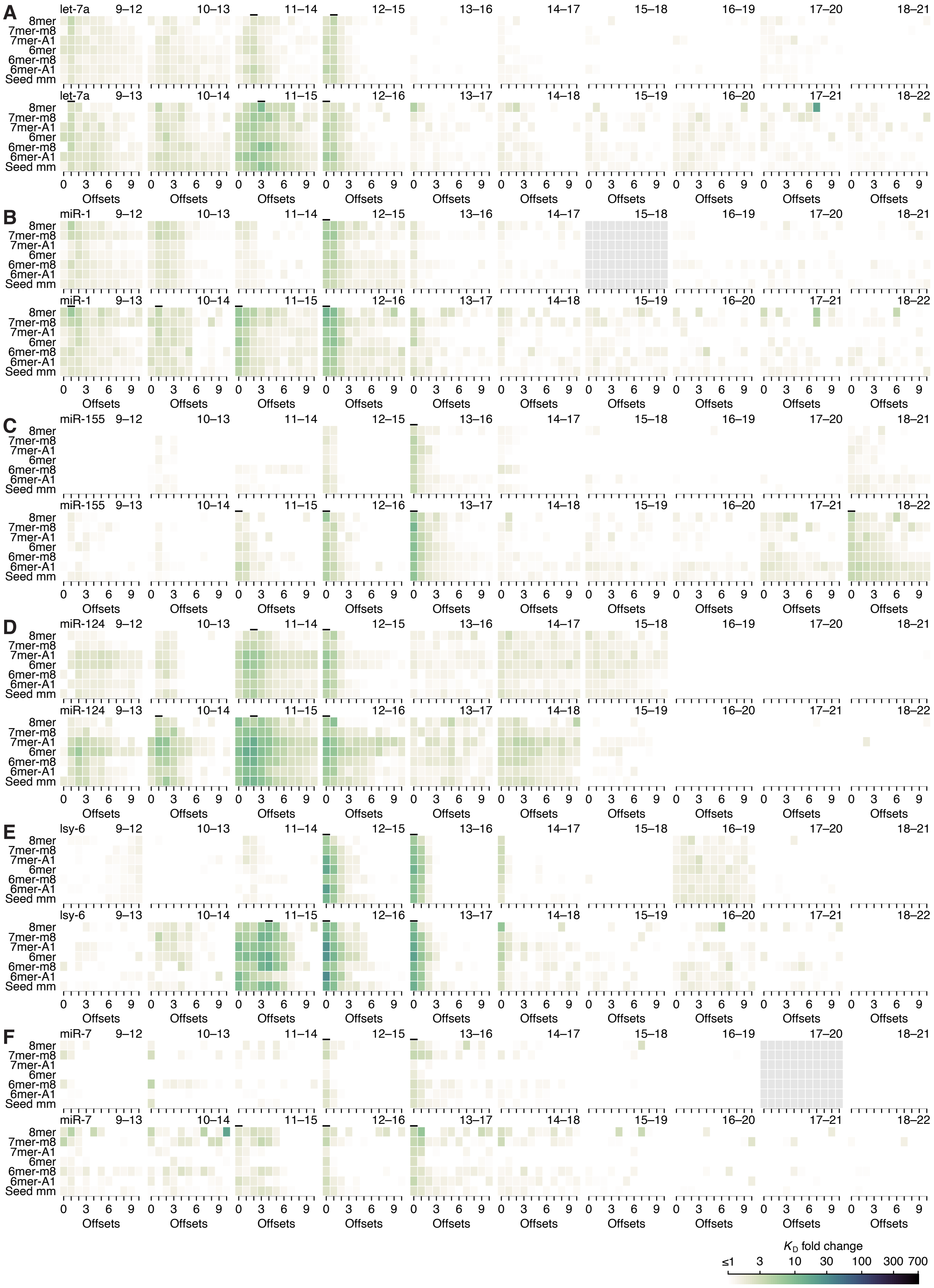
A
B
C
D
E
F
let-7a
miR-1
miR-155
miR-124
lsy-6
miR-7
9–12
10–13
11–14
12–15
13–16
14–17
15–18
16–19
17–20
18–21
9–13
10–14
11–15
12–16
13–17
14–18
15–19
16–20
17–21
18–22
8mer
7mer-m8
7mer-A1
6mer
6mer-m8
6mer-A1
Seed mm
Offsets
0 3 6 9
$K_D$ fold change
≤1 3 10 30 100 300 700

**Figure S11. Minimal influence of seed site type on 3′-supplementary pairing**

(**A**–**F**) Heat maps depicting the $K_D$ fold change of canonical sites and seed-mismatch sites of let-7a (A), miR-1 (B), miR-155 (C), miR-124 (D), lsy-6 (E), and miR-7 (F), using data from previously reported, random-sequence AGO-RBNS experiments. Each individual heat map represents the pairing range to the miRNA 3′ region shown in the upper-right, with each row corresponding either to one the six canonical sites or to a seed-mismatch site given by summing the read counts of all 18 internal 8mer-mismatch sites, and each column corresponding to a different pairing offset ranging from 0 to +10 nt. Those columns with horizontal lines above them were used to compute the average 3′-supplementary pairing for the six canonical sites in the analysis of Figure 4H. $K_D$ fold-change values were not calculated for pairing to positions 15–18 for miR-1 nor for pairing to positions 17–20 for miR-7 because each of these two segments were identical to the 4-nt segment at the very 3′ end of each respective miRNA sequence.

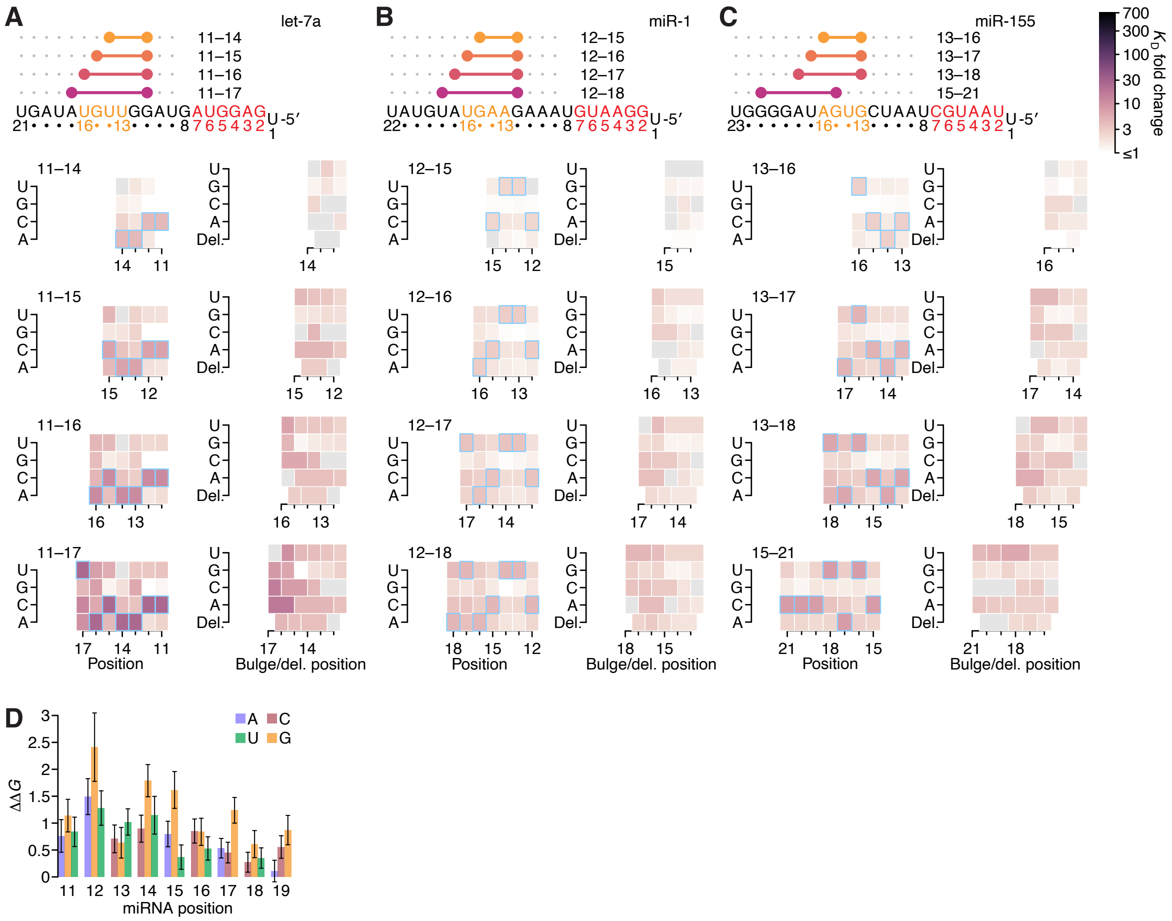

**Figure S12. Further analyses of the impact of mismatched, bulged, and deleted target nucleotides on 3′-compensatory pairing**

(**A**–**C**) The effect of mismatched, bulged, and deleted target nucleotides on 3′-compensatory pairing to let-7a (A), miR-1 (B), and miR-155 (C), in the context of the optimal sites of lengths 4–7 nt for each miRNA. Otherwise, these panels are as in Figure 7A.

(**D**) Profile of mismatch tolerances of sites analogous to those of *C. elegans lin-41*. Each bar represents the Δ$\Delta G$ value of a particular mismatch and position, in the context of a site with contiguous pairing from positions 11 to 19 of let-7a and an offset of +1 nt. Error bars indicate 95% confidence intervals, and colors indicates whether the target nucleotide was changed to an A (blue), U (green), C (red), or G (yellow). A similar polarity in effects, in which mismatches near position 11 are more consequential that those near position 19, is observed when mutating individual *let-7* nucleotides in the animal, although the in vivo experiment observes a more stepwise pattern, in which nucleotides 11, 12, and 13 are each critical for viability, nucleotides 14, 15, and 16 each have intermediate importance, and nucleotides 17, 18, and 19 each have no detectable importance (Duan et al., 2021). This difference might be attributable to the different mismatches examined in the two studies, with our study creating mismatches by changing the nucleotide in the target site and the other study creating a different mismatch by changing the nucleotide in the miRNA. Other possible explanations include differences between human and nematode, and differences between in vitro affinity and in vivo phenotype.

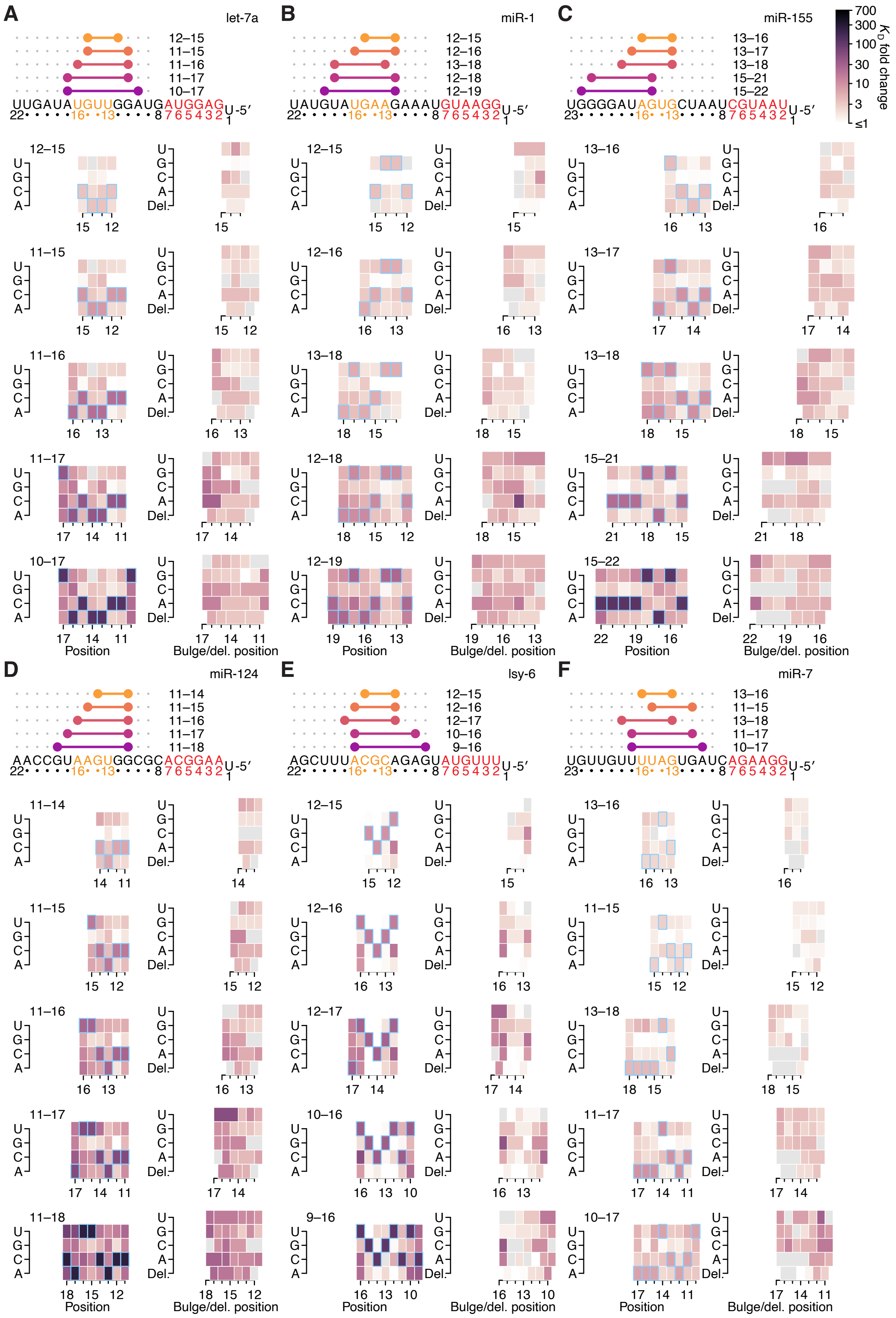
A
let-7a
12–15
11–15
11–16
11–17
10–17
UUGAUAUGUUGGAUGAUGGAGU-5′
B
miR-1
12–15
12–16
13–18
12–18
12–19
UAUGUAUGAAGAAAUGUAAGGU-5′
C
miR-155
13–16
13–17
13–18
15–21
15–22
UGGGGAUAGUGCUAAUCGUAAUU-5′
700
300
100
30
10
3
≤1
$K_D$ fold change
Position
Bulge/del. position
D
miR-124
11–14
11–15
11–16
11–17
11–18
AACCGUAAGUGGCGCACGGAAU-5′
E
lsy-6
12–15
12–16
12–17
10–16
9–16
AGCUUUACGCAGAGUAUGUUUU-5′
F
miR-7
13–16
11–15
13–18
11–17
10–17
UGUUGUUUUAGUGAUCAGAAGGU-5′
Position
Bulge/del. position

**Figure S13. The impact of mismatched, bulged, and deleted target nucleotides on 3′-compensatory pairing in random-sequence AGO-RBNS experiments**

(**A**–**F**) The effect of mismatched, bulged, and deleted target nucleotides on 3′-compensatory pairing to let-7a (A), miR-1 (B), and miR-155 (C), miR-124 (D), lsy-6 (E), and miR-7 (F), in the context of the optimal sites 4–8 nt in length for each miRNA, using data from previously reported random-sequence AGO-RBNS experiments. Otherwise, these panels are as in Figure 7A.
